# S-nitrosylation of EZH2 at C329 and C700 interplay with PRC2 complex assembly, methyltransferase activity, and EZH2 stability to regulate endothelial functions

**DOI:** 10.1101/2024.02.06.579117

**Authors:** Ashima Sakhuja, Yash T Katakia, Shailesh Mani Tripathi, Ritobrata Bhattacharyya, Niyati Pandya Thakkar, S Ramakrishnan, Srinjoy Chakraborty, Sumukh Thakar, Sandeep Sundriyal, Shibasish Chowdhury, Syamantak Majumder

## Abstract

Nitric oxide (NO), a versatile bio-active molecule modulates cellular function through diverse mechanisms including S-nitrosylation of proteins. However, the role of this post-translational modification in regulating epigenetic pathways was very limitedly explored. Herein, we report that NO causes S-nitrosylation of selected cysteine residues of EZH2 in endothelial cells (EC) resulting in SUZ12 dissociation from EZH2 bound PRC2 complex, reduced methyltransferase activity, and diminished nuclear localization eventually hampering its stability. We detected a significant reduction in H3K27me3 upon exposure to NO as contributed by the early dissociation of SUZ12 from the PRC2 complex. Longer exposure to NO donors caused EZH2 cytosolic translocation, its ubiquitination, and further degradation primarily through the autophagosome-lysosome pathway. Through *in silico* S-nitrosylation prediction analysis and site-directed mutagenesis assay, we identified three cysteine residues namely at locations 260, 329, and 700 in EZH2 and further determined that S-nitrosylation of cysteine 329 induced EZH2 instability while S-nitrosylation of cysteine 700 abrogated EZH2’s catalytic activity. A double mutant of EZH2 containing mutations at Cysteine 329 and 700 remained undeterred to NO exposure. Furthermore, reinforcing H3K27me3 in NO exposed EC through the use of an inhibitor of H3K27me3 demethylase, we confirmed a significant contribution of the EZH2-H3K27me3 axis in defining NO-mediated regulation of endothelial gene expression and migration. Molecular dynamics simulation study revealed SUZ12’s inability in efficiently binding to the SAL domain of EZH2 upon S-nitrosylation of C329 and C700. Taken together, our study for the first-time reports that S-nitrosylation dependent regulation of EZH2 and its associated PRC2 complex influences endothelial homeostasis.

## INTRODUCTION

Polycomb repressive complex 2 (PRC2) is a chromatin-modifying group of proteins that work as repressors of gene expression. PRC2 is formed by three major classes of proteins; the functional enzymatic component of the PRC2, Enhancer of Zeste Homolog-2 (EZH2); suppressor of zeste 12 (SUZ12), a PRC2 subunit and a scaffolding protein, and embryonic ectoderm development (EED) which assembles and stabilizes the PRC2 complex.^1,2^ The importance of EED and SUZ12 in maintaining the HMT activity of EZH2 and in the stability of the PRC2 complex is well studied^3,4^. EZH2 is responsible for methylating the 27^th^ lysine residue of histone H3 (H3K27) which results in the repression of gene expression^2,5^. Many studies have highlighted the importance of EZH2 as a crucial player in mediating cellular differentiation during development and also has a considerable role in various disease conditions^6^. Moreover, EZH2 mediates gene silencing in embryonic stem cells thereby promoting pluripotency, and in contrast also in the activation of differentiation^6,7^. The link between EZH2 regulation via deposition of the methylation marks and cancer progression is well reported describing its role in cancer progression and metastasis^8,9^. Indeed, EZH2 dependent suppression of the tumor suppressor genes during cancer progression promotes tumorigenesis.^1,2^

The role of several post-translational modifications such as ubiquitination^10^, phosphorylation^11^, O-GlcNAcylation^12^, sumoylation^13^, and methylation^14^ in regulating EZH2 and its methyltransferase activity were previously reported. Ubiquitination is known to cause a decrease in the methyltransferase activity of EZH2 leading to its degradation. EZH2 has been shown to be phosphorylated by Akt^3^ and AMPK^4^, which leads to suppression of its methyltransferase activity.^15^ O-GlcNAcylation at serine in the SET domain of EZH2 is associated with regulating its methyltransferase activity^16^. Similarly, other mentioned PTMs at specific residues of EZH2, they can regulate its stability and catalytic activity. However, till date, no studies described how S-nitroyslation of EZH2 regulates its localization, degradation and catalytic activity.

Nitrosylation by S-NO formation is one such PTM that has a potential in regulating protein function, stability, and cellular localization. It occurs through NO, a versatile free radical that forms S-NO by selective modification of the protein at cysteine/methionine residues and mediate numerous biological functions^17^. Moreover, many cells harbors endogenous NO producing machinery including NO synthase class of enzymes. NO synthase (NOS) family of enzymes use L-arginine to endogenously produce NO which plays diverse roles in different cell types^18^. EC is one of such cell type that contain a very cell type specific NO producing machinery named as endothelial NOS (eNOS). eNOS dependent release of NO mediates various signaling cascade which are essential for endothelial migration, survival and growth. Indeed, eNOS driven release of NO uses S-nitrosylation dependent regulation of proteins to govern endothelial functions^19,20,21,22^. However, how eNOS dependent release of NO directs gene expression changes in EC through chromatin regulation remains elusive.

In this study, we addressed whether NO-mediated post-translational modification of histone methyltransferase EZH2 could influence its catalytic activity, localization, and stability. Through the present study, we established that S-nitrosylation of cysteine residue(s) in EZH2 protein amends its function as an epigenetic modulator, thereby proving the role of NO as a direct modulator of epigenetic processes and further regulating gene expression changes. Our study for the first time connects the link between eNOS dependent NO release and its function as epigenetic modulator through regulation of EZH2 to alter chromatin structure and thereby dictate NO dependent gene expression changes.

## RESULTS

### Nitric oxide exposure interplayed with EZH2 including PCR2 assembly, its methyltransferase activity, subcellular localization, and stability

NO is well-reported to regulate gene expression changes in EC^22^. However, whether such an effect is dependent on epigenetic processes specifically via regulation of EZH2 and PRC2 complex was never been reported. We, therefore, assessed the effect of NO exposure on EZH2 protein and its catalytic product H3K27me3 level in EC. NO exposure time-dependently caused a reduction in the level of EZH2 protein level specifically observing a significant reduction at 2 hours post exposure to sodium nitroprusside (SNP, Figs. 1A,B). We also observed a significant reduction in the catalytic product H3K27me3, however, the reduction in H3K27me3 was detected within 1 hour of SNP treatment, much earlier than the degradation of EZH2 (Figs. 1C,D). To confirm if such an effect of NO on EZH2 is exerted through transcriptional regulation, we performed qPCR analysis of EZH2 transcripts which revealed no alteration in EZH2 transcript level upon SNP exposure (Supplementary Fig. 1A).

**Figure 1.**
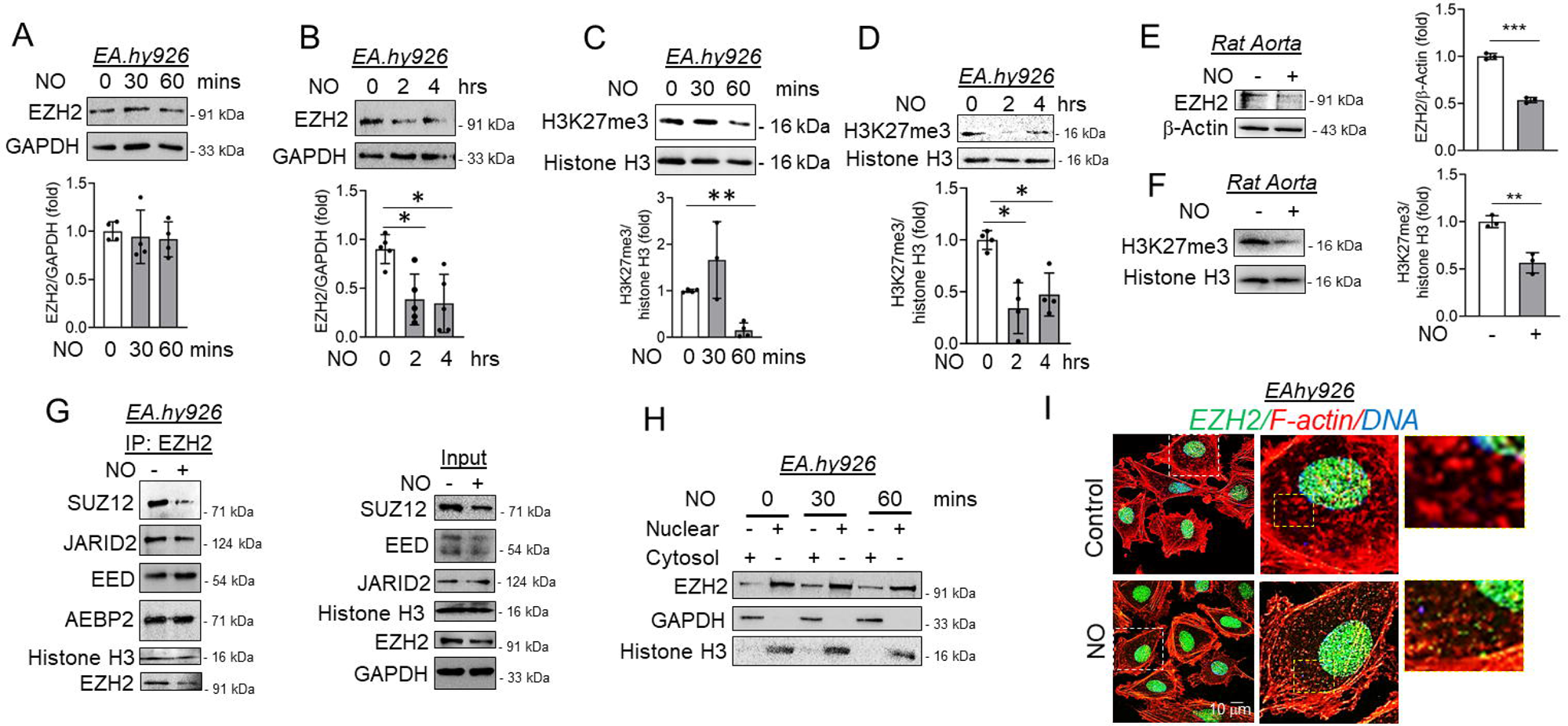
Nitric oxide caused cytosolic localization and degradation of EZH2 protein accompanied with an early reduction in H3K27me3 levels due to dissociation of SUZ12. A-D) Immunoblots analysis of EZH2 (n=4) and H3K27me3 (n=5) proteins levels on exposure to SNP (500 μmol/L) in EA.hy926 cells for variable times. (E-F) Immunoblot analysis using lysate of rat aortic explants exposed to SNP (500 μmol/L) for 2hrs. (n=3) (G) EA.hy926 cells treated with SNP (500 μM) for 30 minutes were subjected to co-immunoprecipitation using EZH2 antibody followed by immunoblotting to show the association of EZH2 with other subunits of the PRC2 complex and histone H3. (n=3) (H) Immunoblotting for EZH2 in nuclear and cytosolic fractions from EA.hy926 cells treated with SNP (500 μmol/L for 30 & 60 mins). GAPDH and histone H3 showed the purity of cytosolic and nuclear fractions respectively. (n=3) (I) Immunofluorescence followed by confocal imaging of EA.hy926 cells exposed to 1 hr of SNP to show cytosolic translocation of EZH2 (green). F-actin (red) is stained with phalloidin-Alexa Fluor 555. DAPI staining is shown in blue (Scale bar: 10 μm). (n=3) Values represent the mean ± SD. *p < 0.05, and **p < 0.01 by unpaired t-test for two groups.

We next confirm such changes in EZH2 and H3K27me3 levels *ex vivo* in rat aorta exposed to NO. Similar to *in vitro* findings, diminished EZH2 and H3K27me3 levels were detected in rat aorta exposed to SNP (Figs. 1E,F). Because EC possesses endogenous NO-producing machinery namely eNOS, we next question whether activation of endogenous NO-production machinery shall also affect the levels of EZH2 and H3K27me3. Induction of EC by bradykinin dose- and time-dependently caused a reduction in EZH2 and H3K27me3 levels (Supplementary Figs. 1B,C). To understand the cell-specific effect of NO on EZH2 and H3K27me3, we next exposed HEK-293, a non-EC type of human cells to NO donor and measured the level of EZH2 and H3K27me3. Through such an experiment, we established that NO exposure caused a reduction in EZH2 and associated H3K27me3 levels independent of the cell type under study suggesting such alteration on EZH2 and its downstream product could be sensitive to NO in many different cell types (Supplementary Fig. 1D). We next evaluated the effect of NO on EZH2 overexpressed through plasmid construct and confirmed that NO exposure also affects the level of overexpressed EZH2 as detected through HA tag or by detecting the total EZH2 (Supplementary Fig. 1E).

Because we observed a reduction in H3K27me3 level much earlier than EZH2 degradation, we next urged whether NO exposure interplays with the assembly of the PRC2 complex which is essential for EZH2 catalytic activity to cause H3K27me3 deposition. We therefore performed a co-immunoprecipitation experiment using EZH2 antibody and detected dissociation of SUZ12 from EZH2 in EC exposed to SNP for only 30 minutes (Fig. 1G). All other components of the PRC2 complex remain associated with EZH2 at least until 30 minutes of NO exposure to EC (Fig. 1G). Interestingly, the protein level of other components of PRC2 including SUZ12, AEBP2, EED, and JARID2 in total cell lysate remain unaltered upon NO exposure to EC (Supplementary Figs. 2A, B).

Being a histone-modifying enzyme and part of the PRC2 complex, EZH2 is primarily localized in the nucleus. However, cytosolic translocation of EZH2 was reported by many earlier studies which also reported its cytosolic substrates including Talin^23^ and small GTPases^24^ to cause actin polymerization. We thus wanted to explore the effect of NO exposure on EZH2 localization. Subcellular fractionation (Fig. 1H) and immunofluorescence (Fig. 1I) followed by confocal imaging strongly indicated cytosolic translocation of EZH2 in EC upon NO exposure. Moreover, confocal imaging also indicated recruitment of EZH2 protein to the filamentous actin (Fig. 1I). No significant changes in localization were detected in other components of the PRC2 including SUZ12, AEBP2, EED, and JARID2 which were primarily localized in the nucleus independent of NO treatment (Supplementary Fig. 2C). Because the turnover of H3K27me3 is not only dependent on the methyl transferase EZH2 but also H3K27me3 specific demethylases UTX and JMJD3, we thus detected the quantity of UTX and JMJD3 protein in EC exposed to NO. UTX and JMJD3 protein levels remained unaltered in EC exposed to NO for 2 hours (Supplementary Figs. 2D,E).

### Nitric oxide exposure caused S-Nitrosylation of EZH2 leading to early SUZ12 dissociation and further altering its binding partners

In the non-canonical NO signaling pathway, many proteins are post-translationally modified through nitrosylation of cysteine/methionine/tyrosine residue thereby regulating the function of such proteins. We, therefore, next questioned whether such could be the case with EZH2 as well. Thus, after establishing the effect of NO exposure on EZH2’s methyltransferase activity, localization, and degradation, we next evaluated whether NO post-translationally modified EZH2 through S-nitrosylation thereby regulating its function. Previous studies reported regulation of EZH2 localization and function via phosphorylation of distinct residues. S-nitrosylation of EZH2 protein has never been reported earlier and therefore we first assessed whether EZH2 is S-nitrosylated upon NO exposure. Through the iodoTMT assay using truncated recombinant EZH2 protein (carrying residues from 429-728 aa) in a cell-free system, we first confirmed that NO exposure caused S-nitrosylation of a truncated form of the EZH2 protein in a cell-free system (Supplementary Fig. 3). Further to confirm this in a cellular system, EC were exposed to SNP followed by Biotin Switch assay and immunoprecipitation using EZH2 antibody. Such an experiment also demonstrated S-nitrosylation of EZH2 protein in EC subjected to NO treatment (Figs. 2A,B). Further to ascertain the S-nitrosylation, we next used a pan S-nitrosylated antibody and confirmed through an immunoprecipitation followed by an immunoblot experiment that EZH2 in EC was indeed S-nitrosylated upon NO exposure (Fig. 2C).

**Figure 2.**
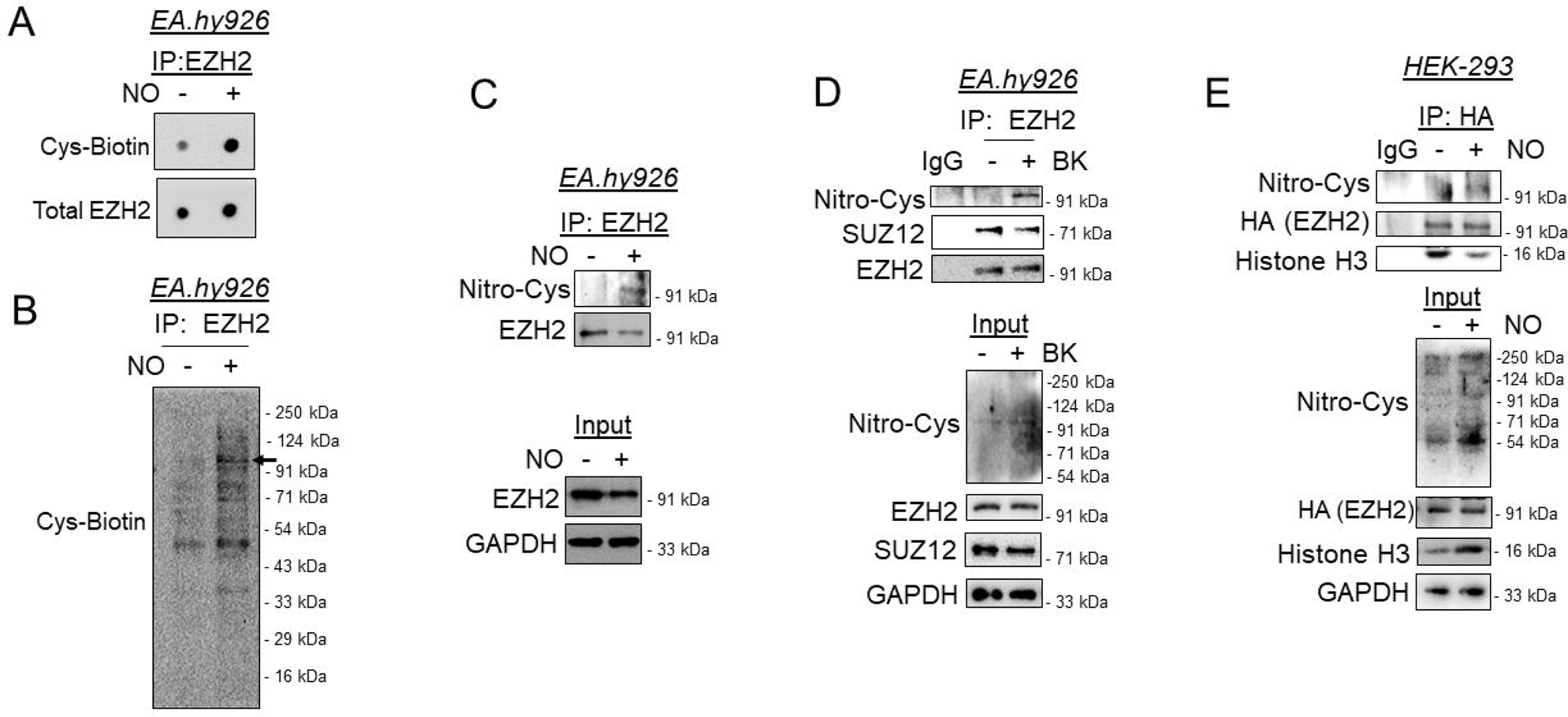
External nitric oxide supplementation or endogenous induction of nitric oxide producing machinery caused S-nitrosylation of EZH2 and dissociation of SUZ12 and histone H3. (A) Dot blot of protein lysates collected from EA.hy926 cells exposed to SNP (500 μmol/L) for 30 min and processed through biotin switch assay followed by immunoprecipitation with EZH2 antibody. Blots were incubated with Streptavidin-HRP followed by developing with chemiluminescence substrate for visualization. (n=3) (B) Same samples were run through SDS-PAGE followed by transfer to nitrocellulose membrane followed by incubation with Streptavidin-HRP and developing the blot with chemiluminescence substrate for visualization. (n=3) (C-D) EA.hy926 cells were exposed to either SNP (500 μmol/L, C) or bradykinin (10 μmol/L, D) for 30 minutes followed by immunoprecipitation with EZH2 antibody and further immunoblotting to show the presence of S-nitrosylation of EZH2 using Nitro-Cysteine antibody. (n=3) (E) Plasmid containing HA tagged EZH2 were transfected in HEK-293 cells followed by exposing to SNP (500 μmol/L) for 30 minutes. Cell lysates were then immunoprecipitated using HA antibody followed by immunoblotting with respective antibodies. (n=3)

We next performed a co-immunoprecipitation experiment using EC which was exposed to bradykinin, a natural inducer of endogenous NO production. In doing so, we detected a robust S-nitrosylation of EZH2 protein concurrent with the loss of EZH2 binding with SUZ12 of the PRC2 complex (Fig. 2D). Furthermore, to support this data, we also analyzed the S-nitrosylation of HA-tagged EZH2 protein in a HEK-293 overexpression system. Through a co-immunoprecipitation experiment using HA antibody, we again confirmed S-nitrosylation of overexpressed EZH2 protein upon exposure to NO (Fig. 2E).

To exclude the possibility of the effect of SNP on EZH2 as non-specific, we next used GSNO which is more well-accepted to impart S-nitrosylation of cellular proteins. In so doing, we found similar reduction in the level of EZH2 after 120 minutes of GSNO exposure (Fig. 3A). As similar to SNP treated EC, further analysis of H3K27me3 in the GSNO treated cells revealed time dependent depletion of H3K27me3 by 60 minutes which remained significantly low at least up to 120 minutes post-GSNO treatment (Fig. 3B). We next wanted to explore the comprehensive interacting partners of EZH2 upon S-nitrosylation to understand the effect of such post-translational modifications on its methylation, translocation and degradation. Mass spectrometric analysis revealed the comprehensive association map of EZH2 in control and GSNO treated cells (Fig. 3C). A total of 261 proteins were identified to be associated with EZH2 while 341 different proteins associated with EZH2 upon GSNO exposure (Figs. 3C, D). Out of 261 proteins in control condition, 48 unique proteins were associated with EZH2 which completely dissociated upon GSNO exposure. In contrast, 128 (out of 341 in total) unique proteins were bound to EZH2 upon GSNO exposure which were not detected to be associated with EZH2 in untreated cells (Figs. 3C-E, Supplementary Tables 1-2).

**Figure 3.**
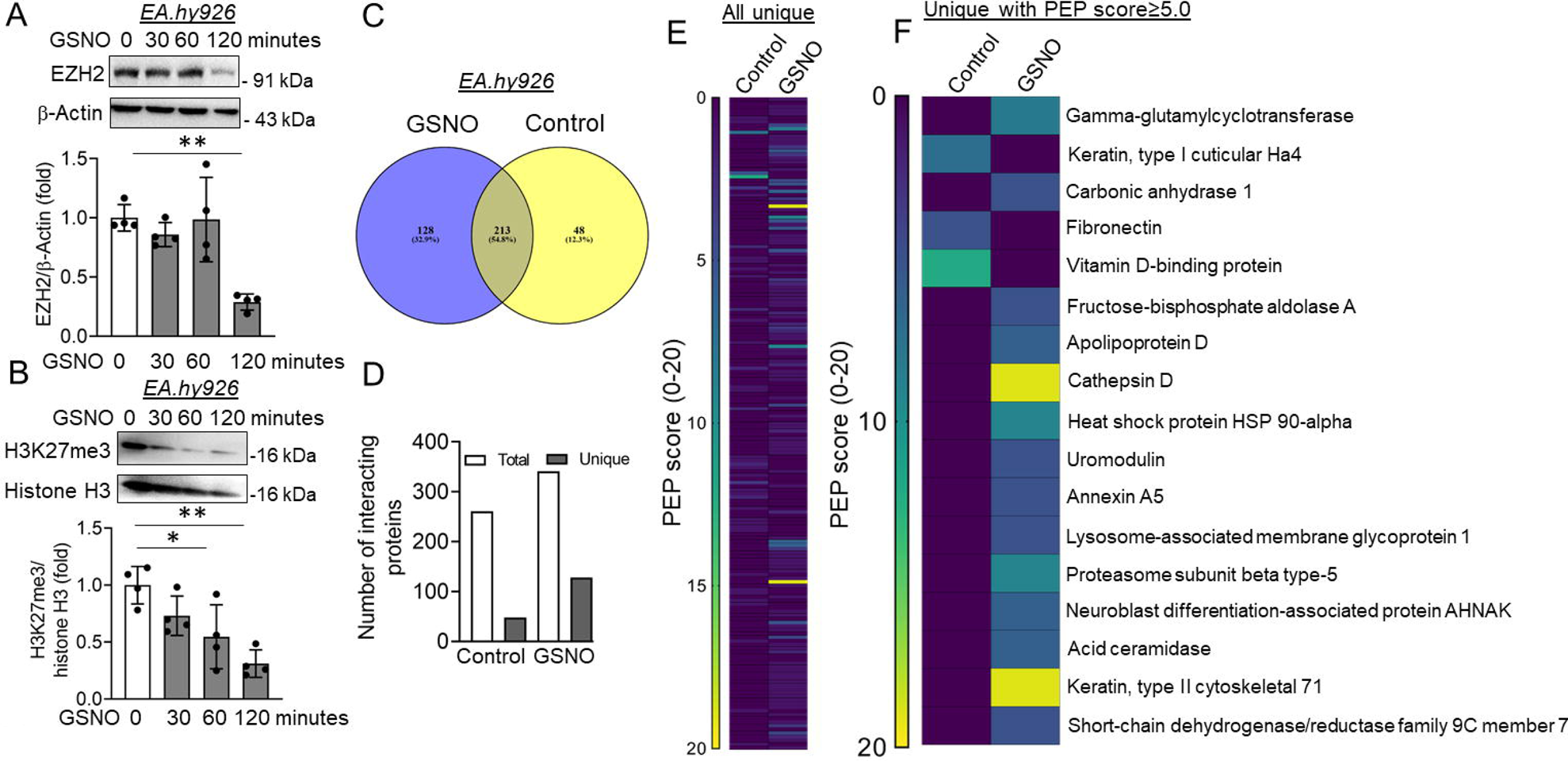
GSNO exposure caused reduction in EZH2 and H3K27me3 level along with altering the interacting partners of EZH2. (A-B) Immunoblotting for EZH2 (A) and H3K27me3 (B) in cultured EA.hy926 cells upon exposure to GSNO (100 μmol/L) for different time points (0, 30, 60, and 120 minutes). (n=4) (C) Venn-diagram show the percentage of common/overlapping proteins in control and 30 minutes GSNO (100 μmol/L) exposed EA.hy926 cells as analyzed through orbitrap mass spectrophotometer. (D) Data showing the number of total and unique interacting proteins present in EA.hy926 cells upon exposure to GSNO (100 μmol/L) for 30 minutes. (E) Heat map based visualization of all the uniquely interacting proteins in control and GSNO treated EA.hy926 cells. (F) Heat map based visualization of uniquely interacting proteins in Control and GSNO treated groups with a PEP score ≥ 5. Values represent the mean ± SD. *p < 0.05, and **p < 0.01 by unpaired t-test for two groups.

We next performed a close analysis of the interactome data which indicated few interesting association patterns of EZH2 in untreated and GSNO treated conditions. Firstly, as observed through co-immunoprecipitation experiment, EZH2 was found to be associated with histone H3 in untreated cells, however, such association was not detected upon GSNO exposure. Surprisingly, nuclear-to-cytosolic shuttling protein 14-3-3 was found to be associated with EZH2 in untreated conditions while a complete loss of association was observed upon GSNO challenge (Fig. 3E, Supplementary Tables 1-2). Although, we expected such association with 14-3-3 to be prominent upon GSNO exposure due to EZH2 cytosolic shuttling after S-nitrosylation, however, we observed an opposite correlation. Such data indicated that EZH2 may be using 14-3-3 during natural cytosolic localization while its cytosolic localization upon S-nitrosylation is likely to be driven by other unknown factors. More interestingly, upon GSNO exposure, we detected EZH2 association with many proteins of the endosome/lysosome/proteasome pathway proteins including several Rab family of proteins, HSP70&90 chaperon proteins, cathepsin D lysosomal protease, lysosome-associated membrane glycoprotein 1, proteasome subunit beta type-5/alpha type-4, lysosomal acid ceramidase (Figs. 3E-F, Supplementary Tables 1-2). EZH2 was not found to be associated with these proteins in control conditions.

### Nitric oxide caused the degradation of EZH2 primarily through autophagosome-lysosome pathway while inhibition of endogenous nitric oxide machinery reversed nitric oxide dependent degradation, activity and localization of EZH2

Because we observed association of many endosomal/lysosomal and proteasomal proteins, we therefore questioned the role of lysosomal and proteasomal degradation pathways in S-nitrosylation dependent degradation of EZH2. The role of post-translational modification-dependent regulation of EZH2 protein degradation is least studied. Because we observed EZH2 protein degradation upon NO exposure, we therefore explored the key degradation machinery responsible for S-nitrosylated EZH2. Because ubiquitination of proteins plays a key role in degradation and previous report indicated ubiquitination mediated degradation of EZH2^25^, we first assessed the level of EZH2 ubiquitination upon NO exposure and found that S-nitrosylated EZH2 are heavily ubiquitinated upon NO exposure (Fig. 4A). Cells use lysosomal and proteasomal degradation pathways for protein degradation, we thus used pharmacological inhibitors of proteasomal and autophagosome-lysosome pathway to evaluate their relative contribution towards degradation of S-nitrosylated EZH2. Inhibition of proteasomal pathway using MG132 was unable to reverse the level of EZH2 upon NO exposure (Fig. 4B). In contrast, inhibition of autophagosome-lysosome pathway using bafilomycin A significantly although partially reversed EZH2 degradation upon NO challenge (Fig. 4C). We then inhibited both proteasomal and autophagosome-lysosome pathway to evaluate the effect of such combination inhibition on EZH2 protein level. In so doing, we detected complete reversal of EZH2 protein level upon combination inhibition of proteasomal and autophagosome-lysosomal pathways (Fig. 4D). All these data indicated that degradation of EZH2 upon NO exposure primarily occurs through autophagosome-lysosome pathway, however, an inhibition of the autophagosome-lysosome pathway could likely switch S-nitrosylated EZH2 degradation through proteasomal pathway. Successful inhibition of autophagosome-lysosomal pathway upon bafilomycin A treatment was confirmed through increase in p62 level (Fig. 4E).

**Figure 4.**
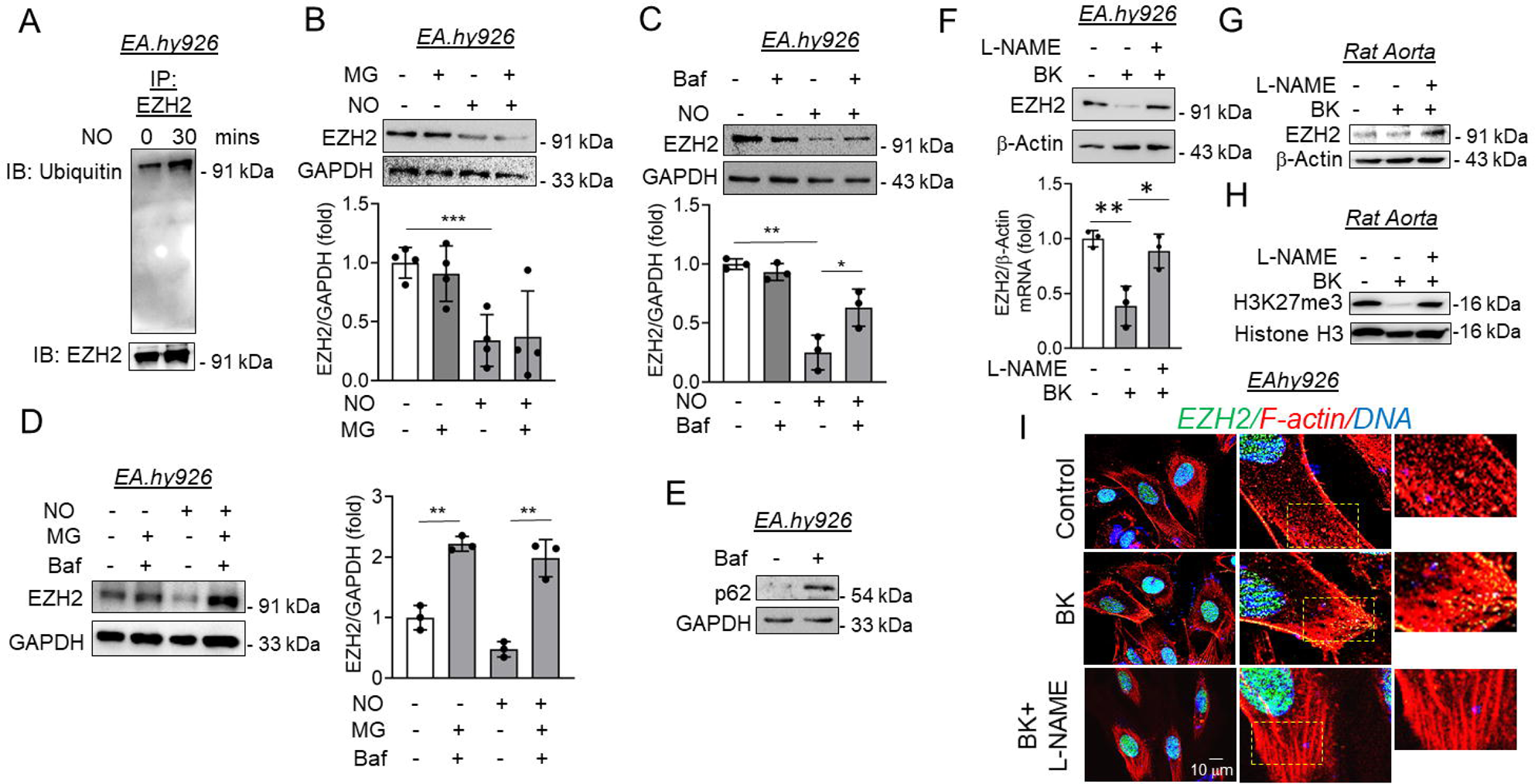
Elevated ubiquitination of EZH2 on exposure to Nitric oxide in endothelial cells to cause its degradation primarily through the autophagosome-lysosome pathway. (A) EA.hy926 cells exposed to SNP (500 μmol/L) for 30 minutes followed by co-immunoprecipitation to pulldown EZH2. This pulldown samples were then subjected to immunoblotting with ubiquitin antibody to show the presence of ubiquitinated EZH2. (B) Immunoblotting analysis of EZH2 in EA.hy926 cells pretreated with MG-132(MG) (1 μmol/L) for 2 hrs followed by exposing to SNP (500 μmol/L) for another 2 hrs. (n = 4) (C) Immunoblotting for EZH2 in cultured EA.hy926 cells pretreated with Bafilomycin A1(Baf) (100 nmol/L) for 2 hours followed by exposing to SNP (500 μmol/L) for additional 2 hrs. (n=4) (D) Immunoblotting for EZH2 protein in cultured EA.hy926 cells pretreated with both MG-132 (1 μmol/L) and bafilomycinA1 (100 nmol/L) for 2 hrs followed by exposing to SNP (500 μmol/L) for additional 2 hrs. (n=4) (E) Immunoblotting to show the presence of elevated p62 in EA.hy926 cells upon treatment with bafilomycin A1 (100 nmol/L) to confirm disruption of autophagosome-lysosome pathway. NOS inhibitor treatment reversed NO dependent reduction in EZH2 and H3K27me3 level. (F) Immunoblotting for EZH2 with protein lysate collected from cultured EA.hy926 cells pretreated with L-NAME (1 mmol/L) for 1 hr followed by exposing to bradykinin(BK) (10 μmol/L) for additional 2 hrs. (n=3) (G-H) Rat aortic rings were pretreated with L-NAME (1 mmol/L) for 1 hr followed by exposing to bradykinin (10 μmol/L) for additional 2 hrs. Immunoblot analysis were carried out for EZH2 (G) and H2K27me3 (H) in these tissue lysates. (n=3) (I) EZH2 (green) immunostaining along with FLactin (Red) staining of EA.hy926 cells after pretreating with L-NAME (1 mmol/L) for 1 hr followed by exposing to bradykinin (10 μmol/L) for additional 2hrs. DAPI staining is shown in blue. (Scale bar: 10 μm) (n=3) Values represent the mean ± SD. *p < 0.05, and **p < 0.01 by unpaired t-test for two groups.

We next questioned whether inhibition of endogenous NO producing machinery could alter the downstream effect of natural inducers of endogenous NO production machinery in EC such as bradykinin. We exposed the EC with L-NAME, a nonselective inhibitor of all type of NO synthase (NOS) prior to inducing with bradykinin. As observed earlier in our study, bradykinin induction caused reduction in EZH2 level which is completely reversed upon inhibition of NOS with L-NAME (Fig. 4F). We also performed the assay in *ex vivo* rat aorta model which further revealed reversal of EZH2 protein degradation and protection of H3K27me3 in aortic tissues exposed to L-NAME prior to bradykinin treatment (Figs. 4G,H). We then performed localization analysis of EZH2 in cells exposed to bradykinin alone or in combination with L-NAME. As expected, bradykinin induction caused cytosolic translocation of EZH2 which are also found to be localized with actin cytoskeleton, however, inhibition of NOS family of protein in bradykinin treated EC using L-NAME restricted EZH2 localization to nucleus (Fig. 4I).

### Protecting the level of EZH2 downstream product H3K27me3 through inhibition of demethylases reversed nitric oxide dependent effect on endothelial migration and gene expression changes

EZH2’s effect on cellular function is primarily dependent on its regulation of gene expression changes through repressive H3K27me3 mark in the chromatin. Moreover, NO signaling pathway converges to changes in expression of genes associated with endothelial survival, proliferation and migration. We therefore wanted to investigate whether EZH2 dependent catalysis of H3K27me3 play any role in dictating NO driven regulation of endothelial function and gene expression changes. To do so, we took a retrograde approach in which we pre-incubated the cells with GSK-J4, a selective inhibitor of H3K27me3 specific demethylase JMJD3 and UTX. We first confirm that inhibition of JMJD3 and UTX reversed NO dependent reduction in H3K27me3 level (Fig. 5A). We next performed endothelial migration using wound healing assay and observed that NO induced endothelial migration is abrogated upon preserving the level of H3K27me3 through inhibition of demethylase JMJD3 and UTX (Fig. 5B).

**Figure 5.**
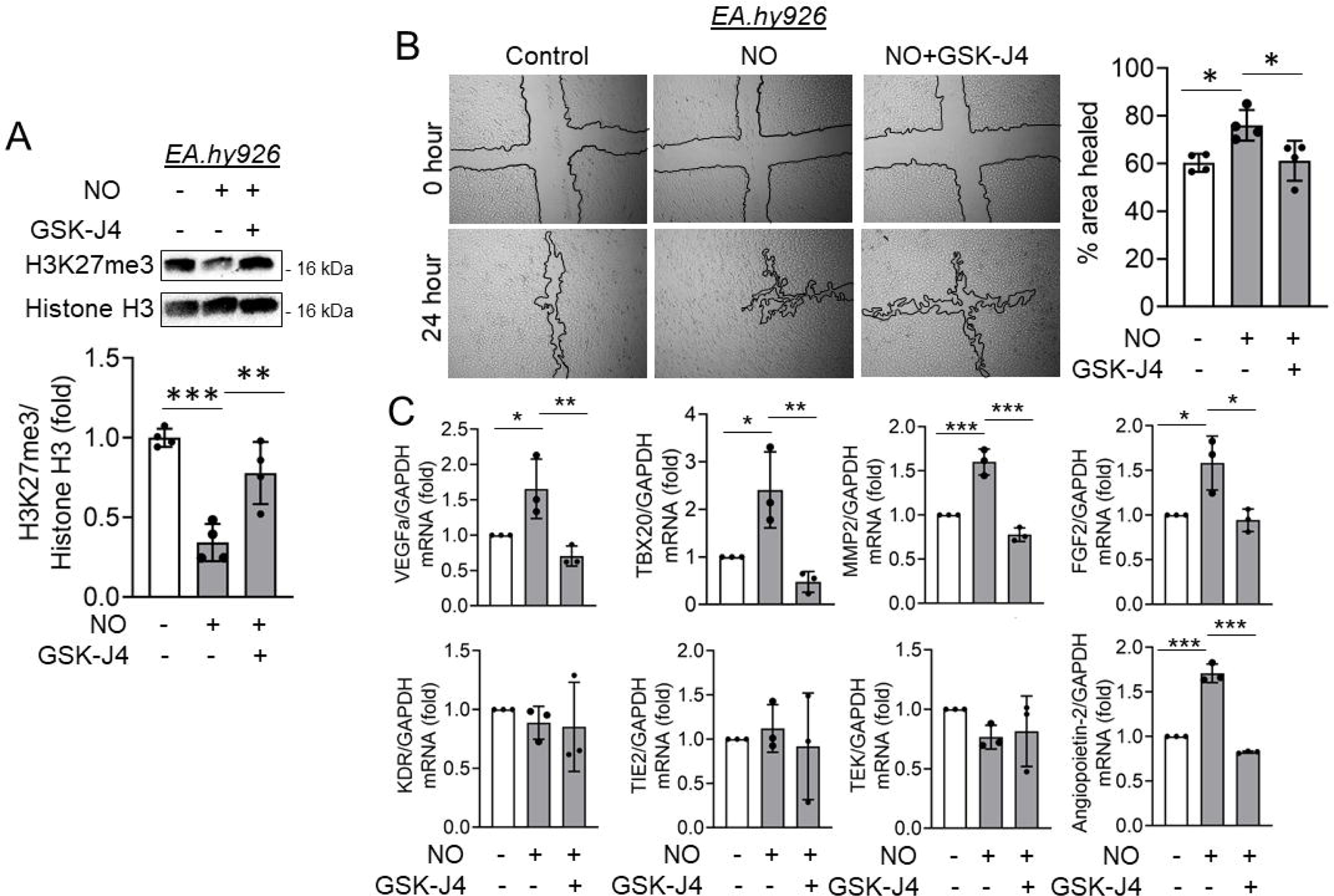
GSK-J4, a demethylase inhibitor protects H3K27me3 level in NO exposed endothelial cells and reversed NO-dependent cell migration and gene expression changes. (A) Immunoblotting for H3K27me3 in EA.hy926 cells pretreated with GSK-J4 (5 μmol/L) for 4 hrs followed by challenging them with SNP (500 μmol/L) for additional 2 hours (n=4). (B) Scratch wound healing assay to show the healing rate of EA.hy926 cells which are pretreated with GSK-J4 (5 μmol/L) for 4 hrs followed by treatment with SNP (500 μmol/L). Healing rate was followed until 24 hrs post-SNP treatment. Images were acquired using bright field microscope adapted with a camera for phase contrast imaging. (n=4) (C) RT-qPCR analysis to measure the transcript level expression of VEGFa, TBX20, MMP2, FGF2, KDR, TIE2, TEK, and Angiopoietin in EA.hy926 cells pretreated with GSK-J4 (5 μmol/L) for 4 hrs followed by challenging them with SNP (500 μmol/L) for additional 4 hrs (n=4). Values represent the mean ± SD. *p < 0.05, **p < 0.01, and ***p < 0.001 by unpaired t-test for two groups.

In similar experimental settings, we next detected the transcript level expression of NO-responsive genes in EC using qPCR experiment. Based on previously reported data sets, we choose to detect the transcript level of NO responsive genes; VEGFa, TBX20, MMP2, FGF2, KDR, TIE2, TEK, and Angiopoietin-2^26^. Through this analysis, we detected significant increase in expression of VEGFa, TBX20, MMP2, FGF2, and Angiopoietin-2 upon NO exposure (Fig. 5C). Pretreatment of EC with GSK-J4 prior to NO exposure completely abrogated NO dependent increment in VEGFa, TBX20, MMP2, FGF2, and Angiopoietin-2 transcript level (Fig. 5C). Surprisingly, although reported earlier to be responsive to NO in EC, in the present experimental settings, we were unable to record any changes in transcript level of KDR, TIE2, and TEK genes (Fig. 5C).

### S-nitrosylation of EZH2 at cysteine 329 and cysteine 700 is important for its stability and methyltransferase activity respectively

Upon confirming the effect of NO on S-nitrosylation of EZH2 and associated changes in its stability, catalytic activity, PRC2 assembly and translocation, we next focused on detecting the possible residues of EZH2 that could be S-nitrosylated. To confirm this, we used GPS-SNO (http://sno.biocuckoo.org/) prediction tools as stipulated in methodology section. Such analysis predicted three possible sites at cysteine 260, 329, and 700 of EZH2 protein with score of 3.158, 2.576, and 3.109 respectively which is beyond the set cut off value of 2.443 (Supplementary Fig. 4A). Predicted cysteine residues 260 and 329 lies within the domain II of EZH2 which essentially allows SUZ12 association with EZH2, in contrast, predicted cysteine residue at 700 lies within the catalytic SET domain of EZH2 which is essential for its enzymatic activity.

To further decipher the role of each of these cysteine residues, we generated point mutated constructs of these cysteine by using site-directed mutagenesis kit to convert the codon to code for serine in places of the actual cysteine (TGT/TGC to AGT/AGC) residues (Supplementary Fig. 4B). Using overexpression of these constructs in HEK-293 cells, we first performed biotin switch assay to assess the S-nitrosylation of these mutated EZH2. Such analysis revealed partial loss of S-nitrosylation of EZH2 for each of the mutants EZH2 C260S, EZH2 C329S, and EZH2 C700S. Interestingly, a complete loss of S-nitrosylation signal was not detected with single mutation indicating multiple cysteine residues were likely to be S-nitrosylated in EZH2 protein upon NO exposure (Fig. 6A). We then analyzed the effect of NO exposure on EZH2 protein and its catalytic product H3K27me3 level in cells overexpressed with WT and mutated form of the EZH2 gene. As observed earlier, NO exposure caused significant loss of HA tagged EZH2 protein and H3K27me3 level in cells overexpressed with HA tagged EZH2 WT gene (Figs. 6B,C). A comparable loss of HA tagged EZH2 protein and H3K27me3 level was also detected in cells overexpressed with HA tagged EZH2 C260S mutant gene (Figs. 6B,C). Interestingly, NO exposure did not alter the level of HA tagged EZH2 protein in cells overexpressed with HA tagged EZH2 C329S mutant gene, however, H3K27me3 level in these cells were significantly reduced upon NO exposure (Figs. 6B,C). In contrast, NO significantly diminished the level of HA tagged EZH2 C700S protein in cells overexpressing HA tagged EZH2 C700S mutant, however, surprisingly, H3K27me3 level upon NO challenge remained relatively unaltered in these cells (Figs. 6B,C).

**Figure 6.**
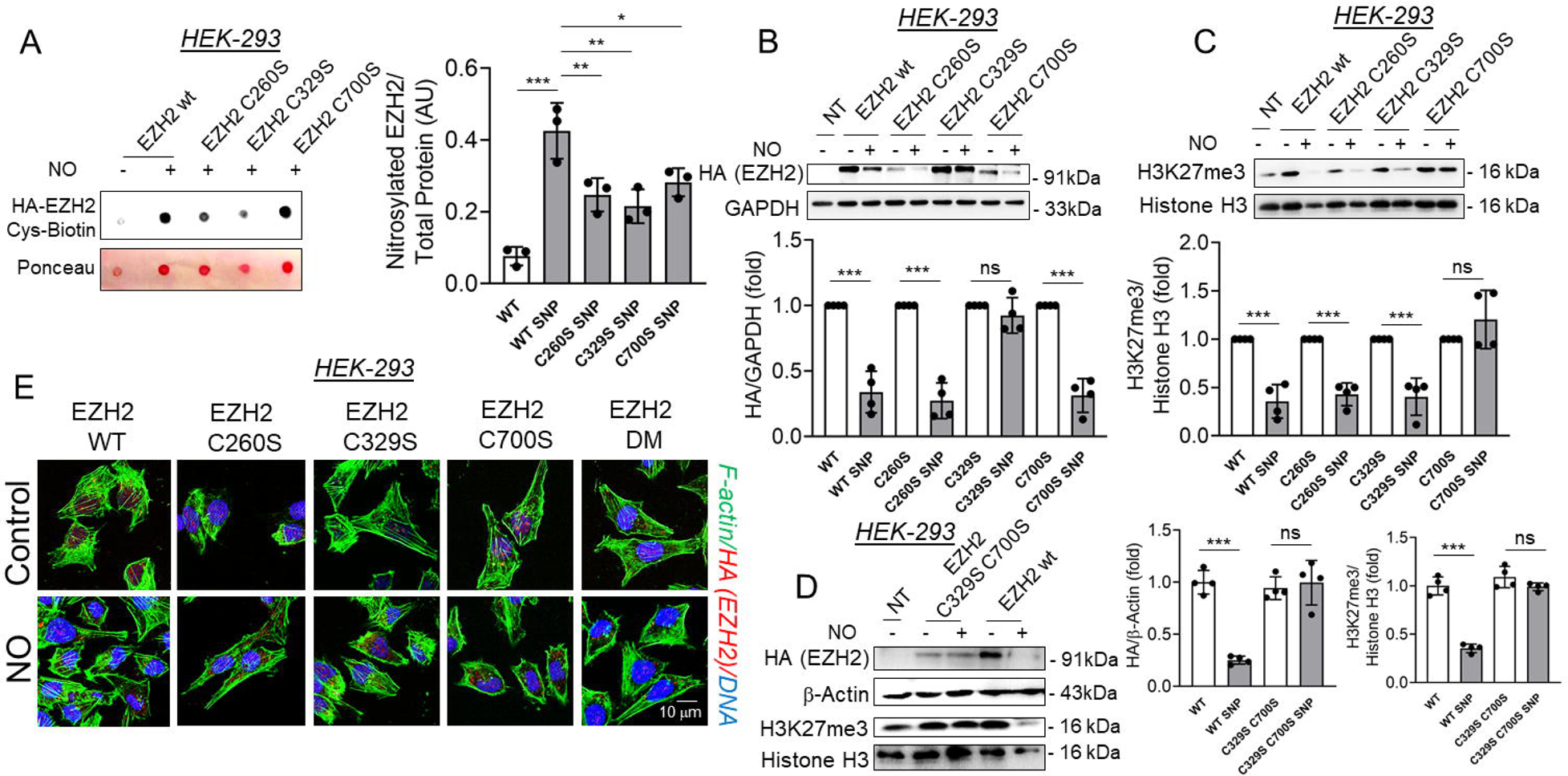
Point mutated EZH2 protein responded differentially to nitric oxide in the context of its effects on the stability and catalytic activity of EZH2. (A) Biotin Switch assay followed by dot-blot and Ponceau staining of HA immunoprecipitated cell lysates of HEK-293 cells overexpressing WT and mutant EZH2 (having point mutations at cysteines 260, 329, and 700) followed by challenging with SNP (500 μmol/L) for 2 hrs to show the levels of S-nitrosylated HA tagged EZH2. (n=3) (B-C) Immunoblotting for HA (EZH2) (B) and H3K27me3 (C) in cell lysates collected from HEK-293 cells containing wild type or mutated versions of EZH2 (C260S, C329S, C700S) and were exposed to SNP (500 μmol/L) for 2 hrs. (n=4) (D) Immunoblotting experiments to show the levels of HA (EZH2) and H3K27me3 in HEK-293 cells transfected with wild-type and the double mutant version of EZH2 (created by inserting point mutations to convert cysteine to serine at positions C329 and C700) and exposed to SNP (500 μmol/L) for 2 hrs. (n=4). (E) Immunostaining for HA (EZH2) (Red) along with FLactin (green) staining in HEK-293 cells to localize HA tagged EZH2 in wild-type and mutated versions, following their exposure to SNP (500 μmol/L) for 2 hrs. DAPI staining is shown in blue. (Scale bar: 10 μm) (n=3) Values represent the mean ± SD. ***p < 0.001 by unpaired t-test for two groups.

We next decided to generate a double mutant form of EZH2 gene where both cysteine residues at location 329 and 700 are mutated to serine thereby generating a double mutant named EZH2 C329S C700S mutant. We then overexpressed this double mutant in HEK-293 cells followed by treating the cells with NO. Double mutated (at cysteine 329 and 700 residues) form of the HA tagged EZH2 was completely insensitive to NO exposure both in context to the level of HA tagged EZH2 C329S C700S protein as well as its product H3K27me3 (Fig. 6D). In similar settings, cells overexpressing WT EZH2 gene showed significant reduction in both HA tagged EZH2 protein and H3K27me3 level (Fig. 6D). We next performed immunofluorescence experiment and confocal imaging study to ascertain the effect of these individual and double mutations of EZH2 on NO driven HA tagged EZH2 localization. In so doing, we observed that NO exposure caused significant cytosolic delocalization of HA tagged EZH2 wt, C260S, and C700S mutated forms of HA tagged EZH2 (Fig. 6E). Interestingly, NO was unable to induce significant cytosolic localization of HA tagged EZH2 C329S and double mutated HA tagged EZH2 C329S C700S protein (Fig. 6E).

### S-nitrosylation of EZH2 protein at C329 and C700 causes conformational changes in EZH2-SUZ12 complex leading to loose association of SUZ12 with the SAL domain of EZH2

Upon experimental validation of EZH2’s S-nitrosylation and its effect on EZH2-SUZ12 interaction, catalytic activity, localization and EZH2 protein stability, we next employed molecular dynamics analysis to understand the effect of S-nitroyslation on EZH2 interaction with SUZ12 protein. Through alphafold modeling of EZH2-SUZ12, we showed the initial conformations of the EZH2-SUZ12 complex in both the EZH2 WT and EZH2 S-nitrosylated at C324 (originally C329) and C695 (originally C700) residues (Fig. 7A) while also clearly visualizing the S-nitrosylation at C324 (Fig. 7B) and C695 (Fig. 7C). The RMSD calculations elucidate the extent to which the conformations of the EZH2 WT and EZH2 S-nitrosylated complexes deviated from their initial states during the course of MD simulations. In comparison to the WT complex, a sharp rise of RMSD values is observed for S-nitrosylated complex after 100 ns indicating possible change in the overall conformations during the simulations. A system showed larger conformational changes for EZH2 S-nitrosylated than for EZH2 WT complex (Fig. 7D).

**Figure 7.**
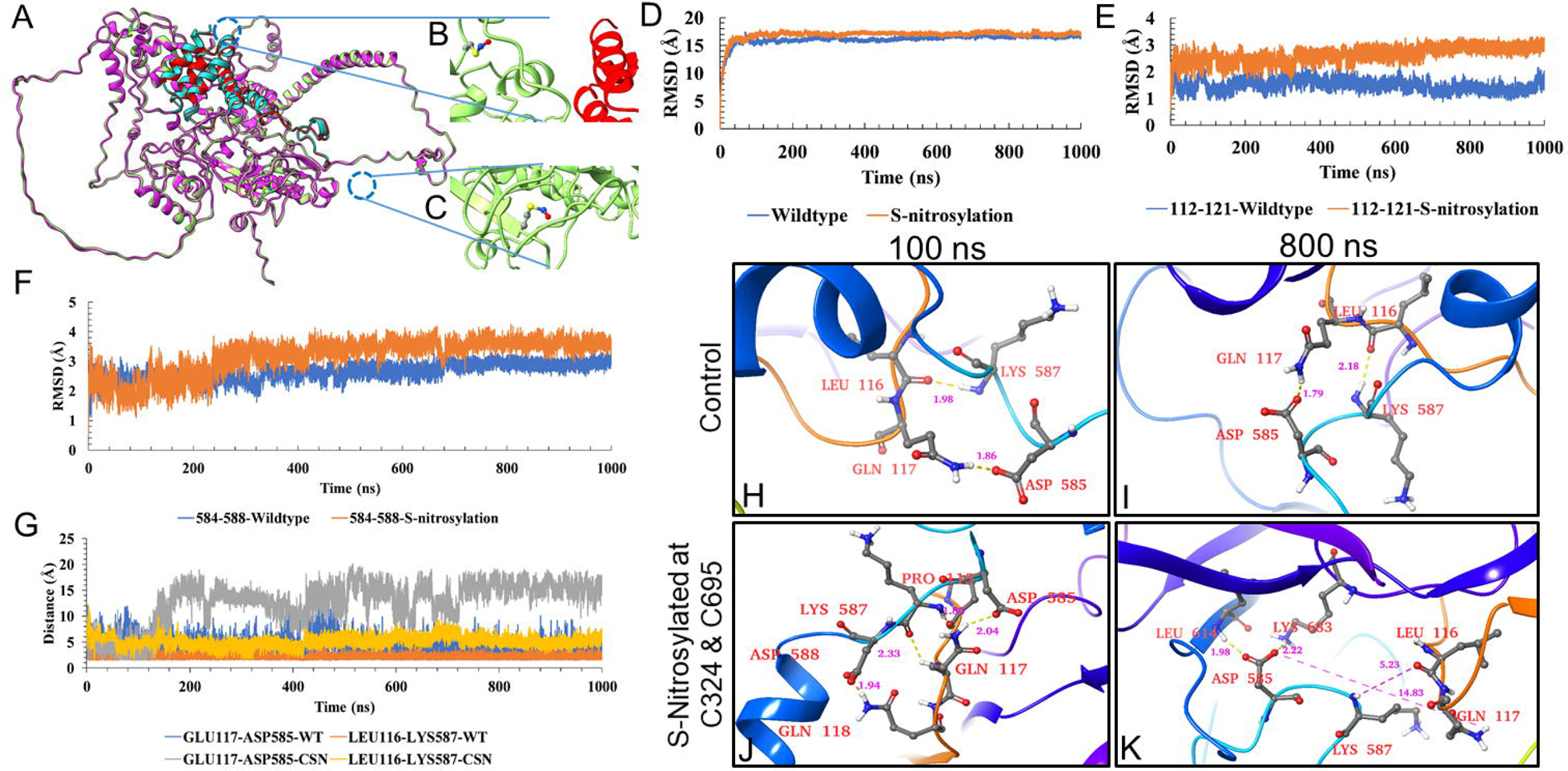
Performing Molecular Dynamic (MD) simulation studies through GROMACS 5.1.5 version using WT and S-nitrosylated EZH2 at C324 and C695. (A-C) Structural illustration of the initial structural alignment of the wildtype and S-nitrosylated EZH2 protein of the EZH2-SUZ12 complex (A). The magenta and green color represents the EZH2 chain of both WT and S-nitrosylated version, while the cyan and red color indicates the SUZ12 chain. It also provides a close-up view of S-nitrosylation at position C324 of EZH2 in the S-nitrosylated protein (B) and also zooms in on the S-nitrosylation at position C695 of EZH2 in the S-nitrosylated protein (C). (D-E) Backbone RMSD of whole EZH2 WT and EZH2 S-nitrosylated (D). Backbone RMSD of EZH2 WT and EZH2 S-nitrosylated specifically within the residue ranges between 112-121 of SAL (E). Backbone RMSD of residues in SUZ12 ranging between 584-588 (F). (H-K) Structural illustration of EZH2-SUZ12 complex near the SAL (112-121) domain of EZH2 in both EZH2 WT and EZH2 S-nitrosylated forms at 100 ns and 800 ns of simulation. Structural data reflects the distance between unique residues of EZH2 (LEU116 and GLN117) and SUZ12 (ASP585 and LYS587) to reflect potential for hydrogen bonds which were lost at 800 ns only in EZH2-SUZ12 complex having S-nitrosylated form of EZH2 at C324 and C695.

Gamblin et al. (2016) reported that residues 112 to 121 within the SAL pack engage with the SET-1 region, serving to stabilize its conformation within the active complex^27^. Importantly for regulation in the human PRC2 complex, the conserved acidic residues 584-588 of SUZ12 reciprocally pack against residues 112-121 of SAL. Consequently, we conducted measurements of the backbone RMSD for both residue ranges, 112-121 of SAL and 584-588 of SUZ12, revealing a significant fluctuation in the RMSD within the SAL region of EZH2 S-nitrosylated with little to negligible RMSD fluctuation for SUZ12 residues ranging from 584-588 (Figs. 7E,F).

Another crucial parameter of MD is root mean square fluctuation (RMSF) to determine the flexibility of protein residues. The RMSF value of both EZH2 WT and EZH2 S-nitrosylated remained relatively stable with a nearly similar trend (Supplementary Fig. 5A). Primarily, the residues near the SAL and long loop region (residue 112-121 and 345-421) display significant fluctuations. Apart from these loops and SAL region of EZH2, RMSF analysis shows stable interactions in both the simulated complexes. Subsequently, we visualized the interface hydrogen bond interactions between SAL residues 112-121 and SUZ12 residues 584-588 throughout 1 µs simulation. Interestingly, in the EZH2 WT, we observed the oxygen atom of LEU166 in EZH2 interacting through the hydrogen bond with LYS587 of SUZ12 and the HE22 atom of GLN117 in EZH2 interacting with the OD2 atom of ASP585 in SUZ12 (Figs. 7H,I) These interactions persisted throughout the simulation. In the case of S-nitrosylated mutant, these two hydrogen bond interactions were initially established and maintained up to 300 ns. However, beyond this time point, the interactions were no longer observed.

Hydrogen bonds (H-bonds) are important non-covalent interactions that help stabilize protein-ligand and protein-protein complexes. To better understand the dynamics of these interactions at the interface, we analyzed the H-bonds formed between specific residues 112-121 of SAL and 584-588 of SUZ12. This examination allowed us to gain insights into the overall stability and binding strength of the complexes (Supplementary Fig. 5B). During the simulations, we closely observed the formation of H-bonds within a distance of 3.5 Å. Upon conducting the analysis, it was discovered that the specific residues 112-121 of SAL were found to interact with 584-588 of SUZ12 in the EZH2 WT complex. This interaction resulted in an average of 2 hydrogen bonds. Conversely, the S-nitrosylated counterpart exhibited a significantly lower average of 0.35 hydrogen bonds suggesting EZH2 WT complex have stronger interactions than EZH2 S-nitrosylated variant. This observation also aligns with the energy profile (Supplementary Fig. 5C). We further measured the distance between the oxygen atom of LEU166 in EZH2 and the hydrogen atom of LYS587 in SUZ12, as well as the distance between GLN117 in EZH2 and ASP585 in SUZ12. A sharp increase in the distance between GLN117 and the interacting residue ASP585 after 150 ns in the EZH2 S-nitrosylation complex simulation (Fig. 7G) was observed, that ultimately led to the disruption of the interaction. Such distance between the oxygen atom of LEU166 in EZH2 and the hydrogen atom of LYS587 in SUZ12, as well as the distance between the HE22 atom of GLN117 in EZH2 and its interacting atom OD2 of ASP585 in SUZ12 at 100 ns (after stabilization of the complex upon initial simulation) and at 800 ns, was also presented through the structure analysis (Figs. 7H-K).

To determine the more accurate free energy binding of all complexes, we utilized the MM/PBSA-based method. The binding free energy in this context refers to the total of all non-bonded interaction energies between the EZH2 and SUZ12 during the MD simulation, including van der Walls, electrostatic, polar solvation, and SASA energies. A more negative binding free energy indicates a stronger affinity between the EZH2 and the SUZ12. We calculated the binding free energies using 1µs MD trajectory, revealing that EZH2 WT and EZH2 S-nitrosylation have a binding free energy of −4649.807 ± 454.449 kJ/mol and −4582.981 ± 453.056 kJ/mol, Subsequently, estimation of their contributions to overall ΔΔG_bind=66 kJ/mol indicates that the contributions vary from one system to the next. On average, the wildtype state tends to have a marginally higher contribution (Supplementary Fig. 5C and Supplementary Table 3). Furthermore, the comparison of the binding energy decomposition between selected residue of SAL and SUZ12 in EZH2 WT and EZH2 S-nitrosylated complexes located at the binding interface reveals that the EZH2 WT predominantly contributes to binding in locally stabilizing manner, Conversely, EZH2 S-nitrosylation displays less contribution to binding energy and these effects might be the consequence of introduced nitrosylation at cysteine 324 and 695 (Supplementary Figs. 5D,E). Overall, these modeling results suggest that S-nitrosylation of the cysteine residues might induce a conformational change in the SAL region which is crucial for the regulation of the PRC2 complex. This conformational change weakens the SUZ12-SAL interaction, ultimately resulting in the destabilization of the SUZ12-EZH2 complex.

## DISCUSSION

A key regulatory pathway of NO-dependent cellular function is through S-nitrosylation of protein which is a reversible post-translational modification. Extensive studies have shown that S-nitrosylation regulates diverse physiological and pathological processes including angiogenesis, adaptive immunity, diabetes, heart failure, stroke etc. through modulating stability, activity, subcellular localization, conformation change, or protein-protein interaction of target proteins^28^. Although NO signaling alters gene expression changes in vascular endothelium^29^, however, the role of NO-dependent S-nitrosylation of proteins in defining the epigenetic landscape of EC is not clearly understood. PRC2 complex is one of the most abundant and well-studied complexes engaged in gene repression through regulation of chromatin compaction. EZH2 being the catalytic unit of PRC2 complex catalyzes the deposition of H3K27me3 mark that causes chromatin compaction and thereby resulting in the repression of gene expression. Nevertheless, how S-nitrosylation regulate EZH2 protein and thereby the activity of the crucial PRC2 complex was never reported. Herein, for the first time we described that NO caused S-nitrosylation of EZH2 protein leading to inhibition of its catalytic activity, disassembly of SUZ12 from PRC2 complex bound EZH2, cytosolic translocation of the protein followed by ubiquitination and degradation through autophagosome-lysosome pathway. Moreover, mass spectrometric analysis revealed S-nitrosylation of EZH2 significantly alters its interactome preferentially allowing association with proteins of the endosome/lysosome/proteasome pathways. We also demonstrated that natural induction of endogenous NO machinery in EC also imparts the exact same effect on EZH2 protein. Through site-directed mutagenesis studies, we specifically identified that S-nitrosylation of Cysteine 329 and Cysteine 700 residues were responsible for EZH2 protein translocation/degradation and catalytic deactivation respectively. Molecular dynamics analysis suggested structural changes in EZH2 protein upon S-nitrosylation causing instability of EZH2-SUZ12 complex. Most importantly, we demonstrated that such S-nitrosylation of EZH2 was responsible for NO-dependent epigenetic regulation of endothelial gene expression and cellular migration (Figure 8).

**Figure 8.**
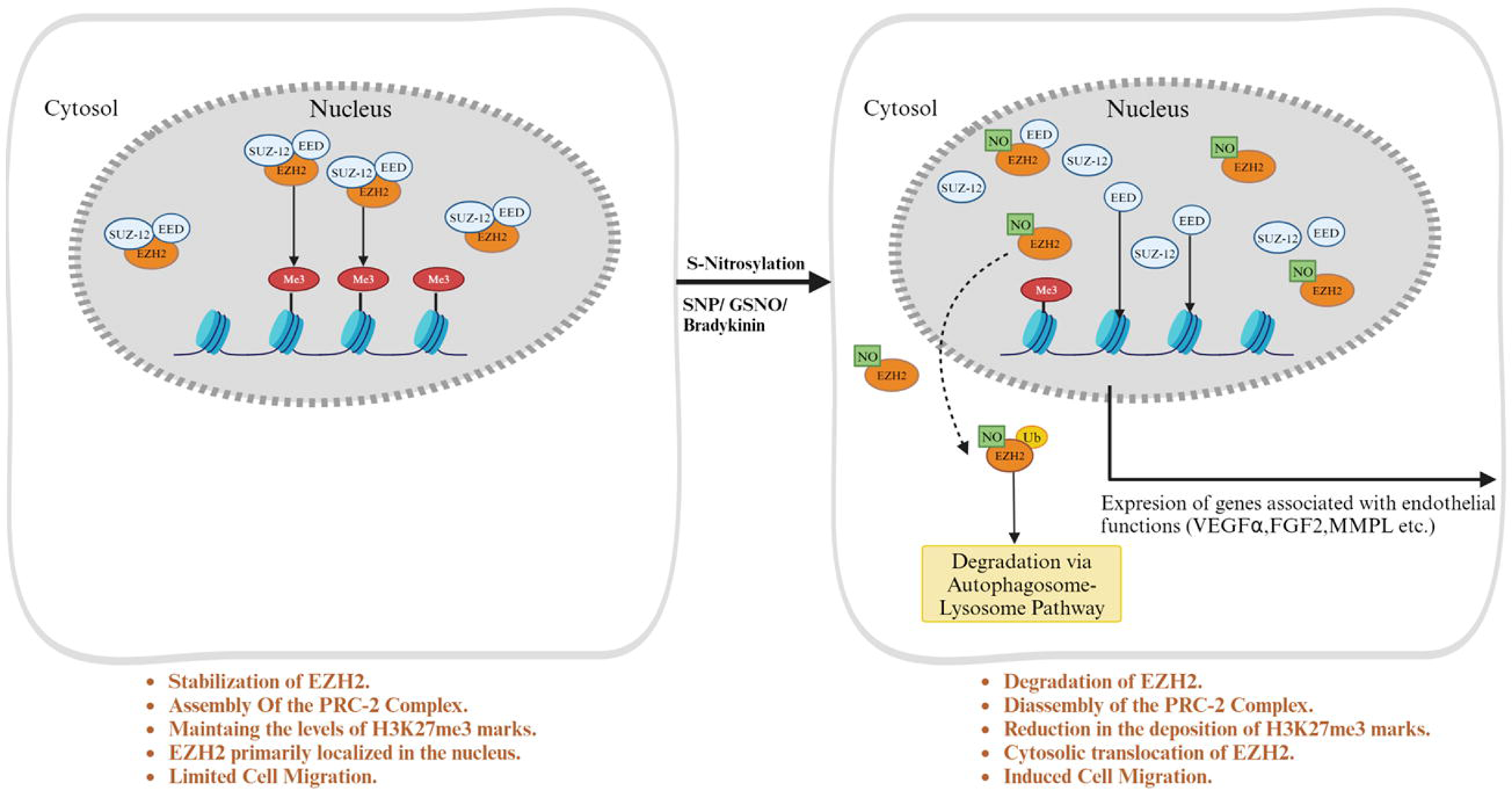
Schematic depicting the effect of EZH2 S-nitrosylation on its stability, translocation and catalytic activity of Polycomb Repressive Complex-2. PRC-2 consists mainly of EZH2, SUZ12, and EED, residing primarily in the nucleus. The major subunit of PRC2 i.e. histone methyltransferase EZH2 localizes primarily in the nucleus and is responsible for deposition and maintenance of the levels of H3K27me3. These cells also have a limited cell migration capacity. On exposure of these cells to NO, S-Nitrosylation of specific cysteine residues on EZH2 occur leading to a decline in their stability which is marked by reduction in EZH2 protein levels along with dissociation of SUZ12 from EZH2 bound PRC2 complex. Upon disassembly, S-Nitrosylated EZH2 translocate from the nucleus to the cytosol where it undergoes ubiquitination followed by its degradation primarily through the autophagosome-lysosome pathway. Additionally, there is a significant effect on the catalytic activity of EZH2, which is shown by reduction in the deposition of the H3K27me3 marks on histone H3. NO dependent S-nitrosylation of EZH2 also causes a spike in the migratory activities of EC due to EZH2 recruitment to the F-actin and likely contributing towards remodeling prior to degradation. The schematic diagram was prepared using BioRender (Agreement number: WB26FEOYBU).

Post-translational modification of EZH2 protein through phosphorylation at several sites were reported earlier^29–32^. Such modifications of EZH2 differentially regulated the localization, function and its association with other proteins of the PRC2 complex. For instance, AMPK phosphorylated EZH2 at T311 to disrupt the interaction between EZH2 and SUZ12 thereby attenuating PRC2-dependent methylation of histone H3 at Lys27 in ovarian and breast cancer. Moreover, such phosphorylation of EZH2 by AMPK caused upregulation of PRC2 target genes, many of which are known tumor suppressors thereby suppressing the growth of tumor cells^32^. Similarly, we observed that S-nitrosylation of EZH2 protein at C329 and C700 caused its dissociation from SUZ12 thereby abrogating the catalytic potential of PRC2 complex. During breast cancer metastases, p38-mediated EZH2 phosphorylation at T367 promotes EZH2 cytoplasmic localization and potentiates EZH2 binding to vinculin and other cytoskeletal regulators of cell migration and invasion. Interestingly, ectopic expression of a phospho-deficient T367A-EZH2 mutant is sufficient to inhibit EZH2 cytoplasmic expression, disrupt binding to cytoskeletal regulators, and reduce EZH2-mediated adhesion, migration, invasion, and development of spontaneous metastasis^29^. In the present study, we observed that S-nitrosylation of EZH2 in specific at C329 caused its cytosolic localization, a transient binding to actin cytoskeleton, ubiquitination of EZH2 protein followed by degradation through autophagosome-lysosome pathway. A very recent study demonstrated DCAF1 mediated phosphorylation of EZH2 at T367 to augment its nuclear stabilization and enzymatic activity in colon cancer cells. Such DCAF1-mediated EZH2 phosphorylation followed by nuclear localization led to elevated levels of H3K27me3 and altered expression of growth regulatory genes in cancer cells^30^. In contrast, herein we report that S-nitrosylation of C329 residue of EZH2 caused its cytosolic localization followed by lysosomal degradation. Moreover, S-nitrosylation at another site C700 led to inactivation of EZH2’s catalytic activity most likely due to early dissociation of SUZ12 protein from PCR2 complex. Although, phosphorylation dependent regulation of EZH2 protein and the PRC2 complex are well documented, however, other post-translational modifications especially NO dependent S-nitroyslation of EZH2 and its regulation of the overall function of this protein has never been reported. Herein, we demonstrated endogenous NO producing machinery of EC as well as external supplementation of NO using donors regulated EZH2 binding with SUZ12, its localization, and protein stability. Such effect of NO on EZH2 contributed towards NO driven gene expression changes and endothelial migration. These are primarily achieved by NO through S-nitroyslation of EZH2 protein at C329 and C700 residues.

In EC endogenous eNOS dependent release of NO drives cell survival, migration and growth typically acting through canonical and non-canonical pathways^33,34^. Canonically, NO induces soluble guanyl cyclase (sGC) and in downstream Protein Kinase G to mediate actin remodeling required for cell migration^35,36^. Moreover, NO signaling also promote endothelial gene expression changes associated with endothelial survival and growth^37,38^. Such gene expression changes by NO in EC was thought to be primarily controlled by gene transcription and mRNA translation via iron-responsive elements^39^. Herein, for the first time, we reported that eNOS dependent NO release instigates non-canonical pathway through S-nitroyslation of EZH2 protein and its product H3K27me3 thereby controlling the epigenetic regulation of gene expression. This was supported by our data wherein preserving the level of EZH2 catalytic product H3K27me3 through inhibition of the demethylases JMJD3-UTX reversed NO dependent gene expression changes in EC. Moreover, we also observed that regulation of EZH2 by NO was independent of the cell type under study which is also been supported through our data in HEK-293 cells exposed to NO. In parallel to well reported function of NO^40,41^ as well as EZH2^42–44^ in actin cytoskeleton remodeling, we observed that NO exposure caused transient localization of EZH2 on the actin cytoskeleton most likely prior to its degradation through autophagosome-lysosomal pathway. In addition, preserving the level of H3K27me3 via inhibition of demethylases reversed NO effect on cell migration. Through the present study, we are unable to establish whether the effect of NO on endothelial migration is mainly driven by its regulation of EZH2-actin cytoskeleton remodeling axis or via regulation of EZH2-H3K27me3-gene expression regulation axis. Nonetheless, NO dependent remodeling of actin cytoskeleton may not alone be driven by canonical sGC-cGMP-PKG-VASP pathway, in addition, regulation of EZH2 via NO may contribute towards its actin remodeling function.

In summary, our study for the first time reported that NO signaling pathway converges to epigenetically regulate gene expression via controlling the stability and catalytic activity of PRC2 complex, in specific the EZH2 component of the complex. This is primarily driven by NO through S-nitrosylation of C329 and C700 residues of EZH2 which promotes early dissociation of SUZ12 from EZH2-PCR2 complex followed by cytosolic localization of EZH2 and its degradation through autophagosome-lysosome pathways. We also showed that such an effect of NO on EZH2 is cell type independent. Moreover, we also reported that NO driven gene expression changes in EC is primarily dependent on NO regulation of EZH2 and its catalytic product H3K27me3. Our molecular dynamics simulation study provided evidence that upon S-nitrosylation of EZH2 at C329 and C700, caused changes in the structure of EZH2 SAL domain leading to dissociation of SUZ12 binding with EZH2. Furthermore, through site directed mutagenesis of EZH2 at C329 and C700 depicted that S-nitrosylation of C329 is responsible for EZH2 cytosolic translocation and degradation while S-nitrosylation of C700 was primarily responsible for catalytic inactivation of EZH2. Our findings shed light into a new mechanism of NO mediated regulation of EZH2 and PCR2 complex to regulate gene expression associated with endothelial function.

## MATERIALS AND METHODS

### Cell culture

Experiments were done primarily in EA. hy926 (immortalized human umbilical vein endothelial cell), purchased from ATCC, Manassas, USA (#CRL-2922). The cells were cultured in Dulbecco’s modified Eagle’s medium (DMEM) (#AL006A; HiMedia Laboratories) supplemented with 10% fetal bovine serum (#RM1112, FBS; HiMedia Laboratories) and 1% penicillin/streptomycin (#10378, PS; Gibco). The cells were passaged every 2-3 days. Human umbilical vein EC (HUVEC) (#CL002-2XT25, HiMedia Laboratories) were cultured and passaged using HiEndoXL™ EC Expansion medium (#AL517, HiMedia Laboratories) along with 4%growth factor, 5% fetal bovine serum and 1% penicillin/streptomycin antibiotic. Human Embryonic Kidney HEK-293 cell line was a kind gift from Prof. Uma Dubey (Department of Biological Sciences, Birla Institute of Technology and Science Pilani, Pilani Campus, India) were maintained in Minimum Essential Media Eagle (MEM) (#AT154; HiMedia Laboratories). All the cells were maintained at 37 ° C with 5% CO2 in a humidified incubator.

### Plasmid and Transfection

Transfection experiments were done with human WT PCMVHA hEZH2 plasmid (#24230, Addgene), a kind gift from Dr. Kristian Helin (Biotech Research and Innovation Centre, University of Copenhagen, Copenhagen, Denmark). HEK-293 cells with 80% of confluency were transfected using 250ng/ml of pCMVHA hEZH2 and Lipofectamine 2000 (#11668030, Invitrogen, Thermo Fisher Scientific) for 4 hrs. After 48 hours of incubation, the cells were then used for immunoblotting and immunofluorescence experiments.

### Inducers and Inhibitors

Sodium Nitroprusside dihydrate (SNP, 500 μM) (#71778, Sigma Aldrich) was used as an exogenous donor of NO in our experiments. Natural induction of NO was activated using 10 μM of Bradykinin (BK) (#B3259; Sigma Aldrich). Inhibition of NO in cell system was achieved by treating the cells with *N*_ω_-Nitro-L-arginine methyl ester hydrochloride (L-NAME, (1 mM) (#N5751; Sigma Aldrich) which is an analog of arginine. Bafilomycin A1 (100 nM) (#11038; Cayman Chemicals) was used as an autophagy Inhibitor. MG132 (1 μM) (#M7449), a proteasomal inhibitor was from Sigma Aldrich. GSK-J4 (5 μM) (#SML-0701), a H3K27me3 demethylase inhibitor was from Sigma Aldrich.

### Prediction of S-nitrosylation site in EZH2 protein using GPS-SNO prediction tools

To identify the cysteine residues that are likely to be S-nitrosylated in EZH2 protein, we performed an *in silico* analysis of the same by using GPS-SNO prediction tools as available at http://sno.biocuckoo.org/. These tools have been developed by a group of scientists by manually collecting 467 experimentally verified S-nitrosylation sites in 302 unique proteins from scientific literature and can be used for prediction of S-nitrosylation sites. The developer performed extensive leave-one-out validation and 4-, 6-, 8-, 10-fold cross-validations techniques to calculate the prediction performance and system robustness^45^. More than 250 publications have used these tools for prediction of possible S-nitrosylation sites. We next used the human EZH2 primary protein sequence available in the addgene for hEZH2 plasmid (#24230, Addgene) and predicted the possible cysteine residues using the GPS-SNO prediction tools. EZH2 in total has 34 cysteine residues, and according to the partially resolved crystal structure of EZH2, none of these residues are involved in di-sulphide bonds and are likely to be cysteine residues with free –SH group. The analysis through the GPS-SNO platform revealed three unique sites at cysteine 260, 329, and 700 which are likely to be S-nitrosylated.

### Subcellular fractionation

Following the treatment with SNP for the given time point conditions, EA. hy926 cells were washed with phosphateLbuffered saline (PBS) followed by scraping. The cell pellet was resuspended in iceLcold PBS containing 0.1% Nonidet PL40 (NPL40) and 1% protease inhibitor (#P8340; Sigma Aldrich) following centrifugation. After 10 mins of centrifugation, the supernatant was collected as the cellular and cytoplasmic fraction, and the nuclear fraction was further pelleted down. The cellular and nuclear lysate obtained was sonicated for 10 s (×2) followed by Immunoblotting experiments.

### Immunoblotting

Cells grown to 80% confluency were provided treatment conditions followed by washing with sterile1xPBS. They were then incubated in RIPA buffer (#9806; Cell Signaling Technology) containing protease inhibitor for lysis after washing with sterile 1X PBS. After scraping, the cells were subjected to repeated cycles of sonication for 10 s (×2). This was followed by a centrifugation step at 10,000 g for 10 min to pellet down the cell debris and collect the supernatant. Protein quantification was done by Bradford assay. After this, the protein samples were mixed with 5× laemmli buffer and heated at 100°C for 10 mins. They were then processed for SDS polyacrylamide gel electrophoresis with 10–250 kDa prestained protein ladder (#PG-PMT2922; Genetix) used for molecular weight reference. Proteins were transferred onto a nitrocellulose membrane (Bio-Rad) at 15 V, 2.5 A for 35 min. The membrane was blocked with 5% skimmed milk for 1 h. The membrane was incubated overnight at 4°C with the primary antibodies: EZH2 mAb (1:1000; #5246), JARID2 Rabbit mAb (1:1000; #13594), SUZ12 Rabbit mAb (1:1000; #3737), EZH1 Rabbit mAb (1:1000; #42088), EED Rabbit mAb (1:1000; #51673), AEBP2 Rabbit mAb (1:1000; #14129). H3K27me3 (1:1000), UTX (1:1000), JMJD3 (1:1000), HALTag Rabbit mAb (1:500; #3724), GAPDH Rabbit mAb (1:2000; #5174), β-actin (1:2000; #3700), H3 Rabbit mAb (1:2000, #4499) (Cell Signaling Technology), S-Nitrocysteine mAb (1:500; #94930) Ubiquitin mAb(1:1000;#3933). Afterward, the blots were washed with TBS-T(x3) and incubated with a secondary peroxidaseLconjugated antirabbit or mouse IgG antibody (1:2000) (#7074 or #7076, Cell Signaling Technology) overnight. These conjugation reactions were detected using the Clarity™ (#1705061) or Clarity™ Max Western ECL Substrates (#1705062) (BioLRad).

### Immunofluorescence and Confocal Microscopy

Following SNP treatment in EA.hy926 & transfected HEK-293 cells were fixed using 2% paraformaldehyde (10 mins). The coverslips were then treated with 0.1% Triton XL100 (5 min) for permeabilizing the cells. Blocking was done with 2% bovine serum albumin (BSA) for 60 mins at room temperature. Afterward, cells were incubated with primary antibodies EZH2 Rabbit mAb (1:1000) or HA-tag (Alexa Fluor™ mAb 647 Conjugate) (1:2000; #37297) overnight at 4°C. The cells were co-stained with Alexa Fluor™ Plus 480 conjugated antirabbit IgG secondary antibody (1:4000; #A32732) (Thermo Fisher Scientific) for 1 h followed with phalloidin for staining F-actin (1:5000; #A22287 or #A12379, Thermo Fisher Scientific) for 30 min. At last, the cells were stained with 1 µM of DAPI (#D9542; Sigma-Aldrich) for 10 min for nuclear staining. Imaging was done using Zeiss ApoTome 2.0 microscope (Carl Zeiss) and fully Spectral Confocal Laser Scanning Microscope (#LSM 880Carl Zeiss).

### Coimmunoprecipitation Experiments

Protein extraction was carried out using 1x RIPA buffer. After washing the Dynabeads (SureBeads™ Protein A Magnetic Beads, #1614013, BioRad) in PBS-T(x3), they were preincubated with the EZH2 mAb, Ha Tagged mAb, ubiquitin mAb or S-Nitrocysteine mAb (1:50) for 2hrs. The beads were then washed and incubated overnight with a protein cell lysate containing 1000 µg of total protein for pulldown of the desired protein from the cell lysate. The cell lysate was then heated at 70° C for 10 mins and magnetized to be separated from the beads. This was then processed for immunoblotting experiments.

### *Ex vivo* experiments using rat aortic tissues

The Institutional Animal Ethics Committee of BITS Pilani, Pilani Campus approved all experimental procedures for the rodent studies. Ex vivo experiments were done using Male Wistar rats aged 12–16 weeks. After giving anesthesia, the rats were dissected from the ventral end. PBS was used to perfuse the heart and aorta, eliminating blood cells from the vessels. The primary aortas were then collected, and the fatty tissue layers were removed carefully. For acquiring the aortic explants, the aorta was cut in a size of 4-5 mm cylindrical pieces. Following a PBS wash, the explants were cultured in HiEndoXL™ EC expansion medium and incubated in the complete growth medium for 12 hrs before initiating the treatment. Afterward, the tissue fractions were homogenized and subjected to sonication after suspending them in RIPA lysis buffer. The protein was estimated using Bradford assay followed by immunoblotting studies.

### RNA Isolation, cDNA Synthesis and qPCR

Reverse transcriptase-quantitative polymerase chain reaction (RT-qPCR) was performed to measure different gene expressions at the transcription level. Upon reaching 80% confluency EA.hy926 cells underwent GSK-J4 treatment for 4 hr followed by SNP exposure for 2hrs. After that, RNA was isolated from the cells using Trizol Reagent (#15596, TRIzol™ Reagent; Life Technologies, Thermo Fisher Scientific). RNA isolation was succeeded by cDNA preparation from 1µg of total RNA using iScript™ cDNA Synthesis Kit (#1708891; Bio-Rad Laboratories, Hercules, CA, United States). The quality and quantity of the RNA was measured using a Nano-Drop spectrophotometer (SimpliNano; GE Lifesciences). Before cDNA synthesis, DNA contamination was removed by pre-treating isolated RNAs with the DNase. This was followed by Real-time PCR where iTaq™ Universal SYBR® Green Supermix (#1725124; Bio-Rad Laboratories) was used with a total master mix volume of 10 μl and GAPDH was taken as the housekeeping gene. Data analysis was done by calculating delta-delta Ct. Details of the primers are as given in Supplementary Table 4.

### Biotin Switch Assay

EA.hy926 cells or EZH2 WT/mutant overexpressing HEK-293 treated with NO donor SNP (500 µM) for 2 hours were processed with the help of a Biotin Switch Assay kit (Abcam) using the standard manufacturer protocol. With this, all the “S-NO” (S-nitrosylated) groups were replaced with Biotin, forming an S-Biotin complex, which was detected by incubation with the streptavidin bound HRP reagent. After this, the samples were immunoprecipitated with EZH2 specific antibody using the above-mentioned protocol. This was followed by dot blot and immunoblotting experiments for the detection of the S-nitrosylation of the EZH2 protein.

### Scratch wound healing Assay

EA.hy926 cells were grown in a 24-well plate. They were pre-treated with GSK-J4 for 4 hrs following SNP treatment at regular intervals(0-24hrs) to measure the wound healing under different treatment conditions or in combination. Imaging was done at definite intervals to calculate the wound healed by cell migration. The wound was created in a straight line. Another perpendicular wound to the first one was drawn to create a cross-shaped wound using a 1mm microtip.

### Site-directed mutagenesis reaction

Phusion Site-Directed Mutagenesis Kit (#F541Thermo Fisher Scientific™) was used to insert point mutations at the specific positions in the pCMV-HA hEZH2 plasmid as mentioned in the manufacturer’s protocol. Prior to this, the primers were phosphorylated using T4 Polynucleotide Kinase (#EK0031, Thermo Fisher Scientific™). After the ligation step, the mutated product was transformed in the competent Dh5-Alpha E. coli cells. The sequence of primers for inserting point mutations at the predicted sites to convert cysteine to serine are provided in Supplementary Table 5.

### Transformation experiments and Plasmid Isolation

The ligation product from the SDM reaction was mixed with 100µl of competent DH5-Alpha cells and kept on ice for 25 mins. This was followed by heat shock at 42° C for 30 seconds. After adding 900 µl of SOC medium, the product was kept at 37° C for 1 hr. This was followed by centrifugation at 3000 rpm for 5 mins. 800 µl of supernatant was removed, and the pellet was resuspended in the remaining 100 µl media. The product was then spread on the LB Agar plate with the help of a spreader and kept overnight at 37° C. A single colony picked from the plate was used to inoculate 5 ml of Luria Broth (LB) containing ampicillin, which was kept overnight at 200 rpm and 37°C. This was used for plasmid isolation, which was performed using the manufacturer’s protocol (#12123, Qiagen Plasmid Minikit). Isolated plasmid was then sent for Sanger Sequencing to confirm the insertion of point mutations at the desired location and then used for further transfection experiments.

### Detection of S-Nitrosylation using iodoTMT labelling ™ Reagent

EZH2 recombinant protein (#MBS2097714, MyBioSource) were treated with GSNO (100μM) for 30 mins. The samples were then processed for iodoTMT labelling with Pierce TM S-Nitrosylation Western Blot kit (#90105, Thermo Fisher Scientific™). A total of 5μg/ml of EZH2 recombinant protein sample was prepared for both control and GSNO treated conditions in 100μl of HENS Buffer. Next, MMTS was added to each, following vortexing for 1 min and incubating it at room temperature for 30 mins to block all free thiols. The protein was then precipitated by adding 600μl of pre-chilled (−20°C) acetone and freezing the samples at −20°C for 1 hr for MMTS removal. The samples were centrifuged at 4°C for 10 minutes (at 12000g). The tubes were inverted to decant the acetone and the white pellet was dried for 10-15 mins. The pellet was resuspended in 100μl of HENS Buffer and 2μl of the iodoTMTzero labeling reagent was added followed by the addition of 4μl of 1M sodium ascorbate and vortexing briefly to mix and incubating it for 2 hrs at room temperature. The samples were then processed for immunoblotting experiments with anti-TMT antibody.

### Sample Preparation for Mass Spectroscopy analysis

EA.hy926 cells were treated with GSNO (100μM) for 30 mins followed by lysis with RIPA buffer. Protein extraction was carried out using 1x RIPA buffer. After washing the Dynabeads (SureBeads™ Protein A Magnetic Beads, in PBS-T(x3), they were pre-incubated with the EZH2 mAb, (1:50) for 2hrs. The beads were then washed and incubated overnight with a protein cell lysate containing 1000ug of total protein for pulldown of EZH2 from the cell lysate. The cell lysate was then heated at 70° C for 10 mins and magnetized to be separated from the beads. Dynabead bound immunoprecipitated proteins were eluted by heating the samples in 1X PBS. Eluted protein per sample was used for digestion and reduced with 5 mM TCEP and further alkylated with 50 mM iodoacetamide and then digested with Trypsin (1:50, Trypsin/lysate ratio) for 16 h at 37 °C. Digests were cleaned using a C18 silica cartridge to remove the salt. Protein per sample was used for digestion and reduced with 5 mM TCEP and further alkylated with 50 mM iodoacetamide and then digested with Trypsin (1:50, Trypsin/lysate ratio) for 16 h at 37 °C. Digests were cleaned using a C18 silica cartridge to remove the salt and dried using a speed vac. The dried pellet was resuspended in buffer A (2% acetonitrile, 0.1% formic acid).

### Mass Spectrometric Analysis of Peptide Mixtures

Experiments were performed on an Easy-nlc-1000 system coupled with an Orbitrap Exploris mass spectrometer. 1µg of peptide sample were loaded on C18 column 15 cm, 3.0μm Acclaim PepMap (Thermo Fisher Scientific) and separated with a 0–40% gradient of buffer B (80% acetonitrile, 0.1% formic acid) at a flow rate of 500 nl/min) and injected for MS analysis. LC gradients were run for 110 minutes. MS1 spectra were acquired in the Orbitrap (Max IT = 60ms, AGQ target = 300%; RF Lens = 70%; R=60K, mass range = 375−1500; Profile data). Dynamic exclusion was employed for 30s excluding all charge states for a given precursor. MS2 spectra were collected for top 20 peptides. MS2 (Max IT= 60ms, R= 15K, AGC target 100%).

### Data Processing

All samples were processed and RAW files generated were analyzed with Proteome Discoverer (v2.5) against the Uniprot Human database. For dual Sequest and Amanda search, the precursor and fragment mass tolerances were set at 10 ppm and 0.02 Da, respectively. The protease used to generate peptides, i.e. enzyme specificity was set for trypsin/P (cleavage at the C terminus of “K/R: unless followed by “P”). Carbamidomethyl on cysteine as fixed modification and oxidation of methionine and N-terminal acetylation were considered as variable modifications for database search. Both peptide spectrum match and protein false discovery rate were set to 0.01 FDR.

### Model and Protein Preparation for *in silico* analysis

The crystal structure of the Human Polycomb Repressive Complex 2 (PDB ID: 5HYN) was acquired from the Protein Data Bank^27^. This complex exhibits multiple missing loops, particularly within the EZH2 residues at positions 1-9, 182-210, 249-256, and 345-421. To address these structural gaps, the EZH2 structure was downloaded from AlphaFold, assigned an AFDB accession code AF-Q15910-F1, and possessed an average pLDDT score of 76.25 (Uniport ID: Q15910)^46^. Subsequently, the EZH2 alphafold model was superposed to the original PDB file (5HYN) to obtain the PRC2 complex. Finally, EED, H3K79M, and JARID2 K116m3 proteins were deleted to keep only the EZH2-SUZ12 complex for further modelling studies. The EZH2-SUZ12 complex was pre-processed using Protein Preparation Workflow tool in Schrödinger suite 2022^47^. Hydrogens were added, charges were assigned, bond orders were refined, and all water molecules and non-standard residues were deleted before proceeding. The protein backbone was minimized by employing OPLS 2005 force field. Further residues Cys324 and Cys695 were S-nitrosylated by Vienna-PTM^48^.

### Molecular dynamics (MD) simulations

The charges of both wild-type and mutant complexes were neutralized by placing a total of 10 Na^+^ ions at positions with high electronegative potential. Both complex and counter ions were then placed in a pre-equilibrated cubic box of SPC/E water molecules. The periodic box of water was extended to a distance of 10 Å from the protein complex and counter ions. Another 442 NaCl molecules were added to the system to maintain a 150 mM salt concentration. The prepared systems were subjected to 50,000 steps of steepest descent energy minimization. The structural fluctuations and stability of the relaxed protein complex were analyzed by time-dependent molecular dynamics (MD) simulation studies using GROMACS 5.1.5. The particle mesh Ewald method (PME) was used for the calculation of electrostatic interactions^49^. Periodic boundary conditions were imposed in all directions. The long-range electrostatic interactions have been calculated without any truncation, while a 10 Å cutoff was applied to Lennard–Jones interactions. The LINCS-like algorithm were employed to restrain hydrogen-containing bonds only and handle long-range electrostatic interactions. SHAKE algorithm was applied to constrain the bond involving hydrogens^50^. The temperature was controlled at 300 K using Langevin dynamics with the collision frequency 1. A time step of 2 fs was used and the structures were saved at every 10 ps interval for the entire duration of the MD run. Equilibrium phase was comprised of two short 1 ns simulations in NVT and NPT ensembles, utilizing a Berendsen thermostat and Parrinello-Rahman barostat respectively at 300K and 1 bar pressure. The unrestrained 1 μs simulation with NPT ensembles at 300 K was considered as production simulation.

### Structure Analysis

Built-in features of GROMACS, such as gmx rms, rmsf, and hbond, were employed to assess the root-mean-square deviation (RMSD), root-mean-square fluctuation (RMSF), and hydrogen bonds (Hbond), respectively, throughout the trajectory. Visualization of the structures was performed using PYMOL and VMD software^51,52^.

### Binding energy calculations using MM/PBSA

The MM-PBSA protocol was employed to determine the effect of PTMs addition on the binding of subunits within a chosen EZH2-SUZ12 complex. The free binding energies of each complex were computed utilizing the equations provided below. Briefly, the provided set of equations were applied to represent an imaginary AB dimer, where A corresponds to EZH2 and B to SUZ12.

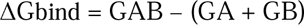

Where GAB is the binding free energy of the EZH2-SUZ12 complex, GA is the binding free energy of the EZH2, and GB is the binding free energy of the SUZ12.

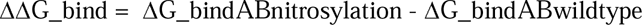

To evaluate the impact of incorporating post-translational modifications (PTMs) into the complex on their binding, we compute the disparity in binding free energy (ΔΔGbind) between the S-nitrosylated and wildtype complexes using Equation 2. The binding free energy was determined by utilizing the 1µs trajectory obtained from molecular dynamics simulations, and calculations were performed on every 500 frames extracted from the last 500 ns of the trajectory using the g_mmpbsa tool integrated with GROMACS^53–55^.

### Statistics Analysis

All the values are expressed as the mean ± SD. An unpaired t-test was used to determine the statistical differences in comparisons between the two groups and one-way ANOVA with a false discovery rate (FDR) method was used for comparison between multiple groups. GraphPad Prism software (GraphPad Prism 9) was used to perform statistical analyses. A p-value (<0.05) was considered statistically significant.

## Data availability

All data can be obtained from the corresponding author on reasonable request.

## Acknowledgment

This work was supported by the Core Research Grant from Science and Engineering Research Board-Department of Science and Technology, Govt. of India (CRG/2022/002209) to SM. This work was also partly supported by a Competitive Research Grant from the Department of Biotechnology, Govt. of India (BT/PR33144/MED/30/2170/2019) to SM. AS is supported by a Senior Research Fellowship from CSIR, Govt. of India (File No-09/719(0107)/2019-EMR-I). Y.T.K., N.P.T. and S.M.T. are supported by a graduate fellowship from BITS Pilani. S.T. was supported by a graduate fellowship from BITS Pilani. S.R. is supported by a graduate fellowship from Department of Science and Technology-Innovation in Science Pursuit for Inspired Research fellowship (DST/INSPIRE/03/2019/000582). We gratefully acknowledge the technical assistance of Mr. Suman Kumar (Confocal Facility, BITS Pilani, Pilani Campus) and BITS Pilani, Pilani Campus for providing the high-performance computing (HPC) facility for simulation study.

## Author contributions

AS designed and performed the experiments, analyzed the data, and wrote the first draft of the manuscript. YTK, RB, NP, SR, SC, and ST performed experiments and analyzed data. SMT, SS, and Shiabsish performed all the molecular simulation studies and wrote the associated methodology and results section of the manuscript for the same. SM secured the funding, designed the experiments, supervised the study and wrote the final draft of the manuscript.

