## Supplementary Figures 1-5 for "S-nitrosylation of EZH2 at C329 and C700 interplay with PRC2 complex assembly, methyltransferase activity, and EZH2 stability to regulate endothelial functions"

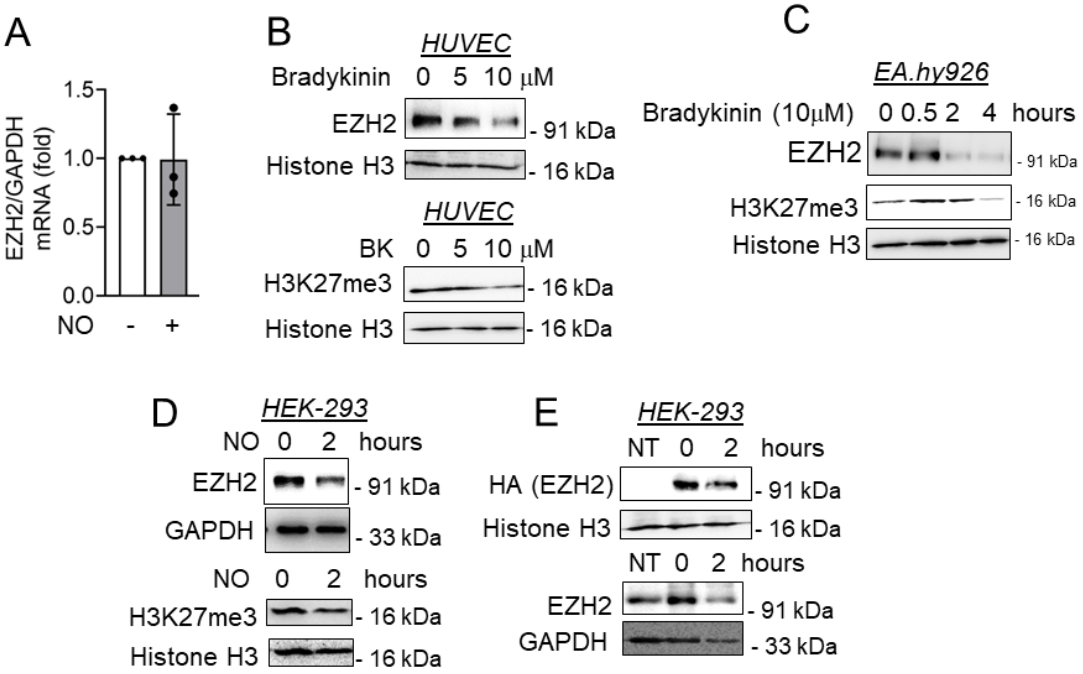


**Supplementary Figure 1. Induction of endogenous nitric or nitric oxide supplementation altered EZH2 and H3K27me3 levels independent of cell types and without altering EZH2 transcript level.** (A) qPCR analysis to measure the transcript level of EZH2 in cultured EA.hy926 cells exposed to SNP (500 μmol/L) for 2 hours. (n=3) (B-C) Immunoblotting for EZH2 and H3K27me3 in HUVEC (B) and EA.hy926 cells (C) exposed to bradykinin(BK) (10 μmol/L). (n=3) (D) Immunoblotting for EZH2 and H3K27me3 in HEK-293 cells on exposure to SNP (500 μmol/L) for 2 hrs. (n=3) (E) Immunoblotting experiment to show the HA-EZH2 and EZH2 levels in the lysates collected from HEK-293 cells transfected with plasmid containing HA tagged EZH2 followed by treatment with SNP (500 μmol/L) for 2 hrs. (n=3)

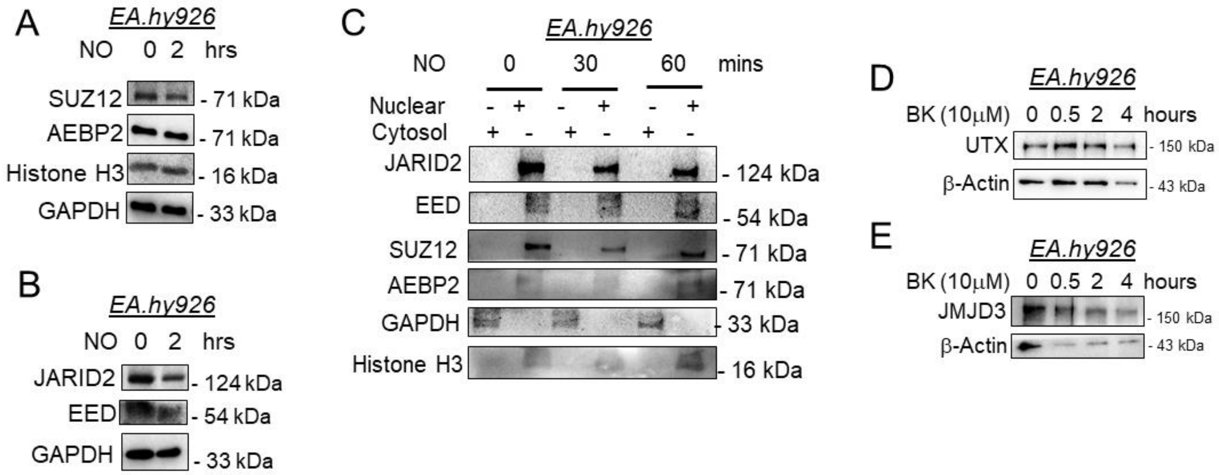


**Supplementary Figure 2. Nitric oxide did not alter the level or localization of other PCR2 complex proteins and no changes were detected in H3K27me3 demethylases UTX and JMJD3.** (A-B) Immunoblotting for SUZ12, AEBP2, and histone H3 (A) along with JARID2 and EED (B) after exposure of EA.hy926 cells to NO donor SNP (500 μmol/L) for 2 hrs. GAPDH was used as controls. (n=3) (C) Sub-cellular fractionation of cell lysates collected from EA.hy926 cells exposed to SNP (500 μmol/L) for 30 and 60 minutes followed by immunoblotting experiment for JARID2, EED, SUZ12, and AEBP2. Presence of GAPDH and histone H3 only in cytosolic and nuclear fractions respectively to indicate the purity of the cellular fractions. (n=3) (D-E) Immunoblotting for UTX (D) and JMJD3 (E) after exposing EA.hy926 cells to endogenous NOS machinery activator BK (10 μmol/L) for different time points. (n=3)

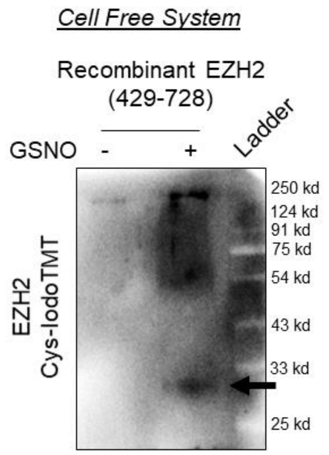


**Supplementary Figure 3. Confirming S-nitrosylation of recombinant human EZH2 (containing aa 429-728) using IdodTMT assay in a cell free system.** In a cell free system, a total of 1 μg recombinant human EZH2 (containing aa 429-728) protein was incubated with GSNO (100 μmol/L) for 30 minutes followed by processing through iodoTMT protocol. Samples were run through SDS-PAGE followed by transferring to nitrocellulose membrane and were incubated with anti-IodoTMT antibody. Blots were developed with chemiluminescence substrate for visualization.

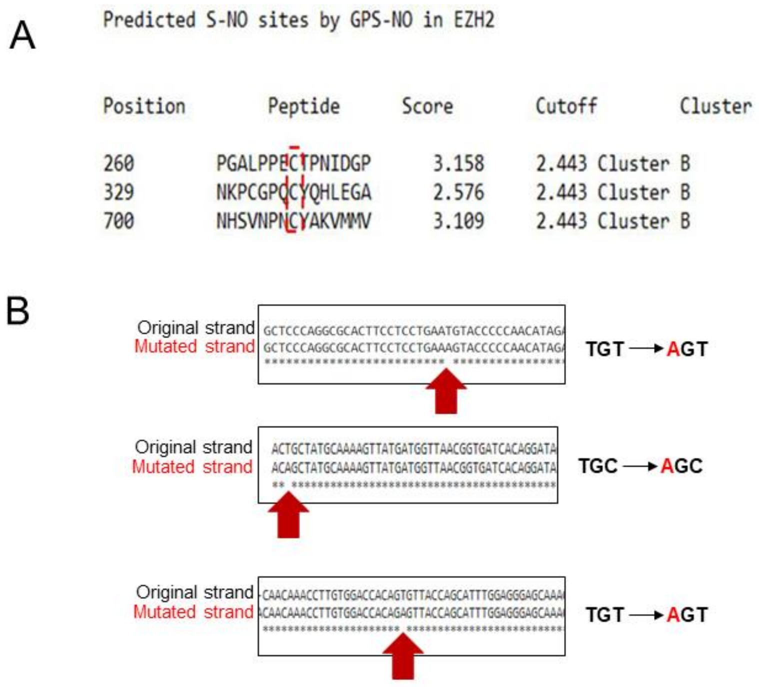


**Supplemental Figure 4. *In silico* S-nitrosylation prediction analysis and generating point mutants for identified residues using site directed mutagenesis kit.** (A) Using pre-validated software, we performed an *in silico* analysis to predict possible cysteine residues that could be S-nitrosylated and identified three cysteine residues at position 260, 329 and 700 (out of 34 total cysteine present in EZH2) that were predicted to be S-nitrosylated. (B) Confirming the insertion of point mutations at these three predicted cysteine sites leading to their conversion to serine using Sanger sequencing technique.

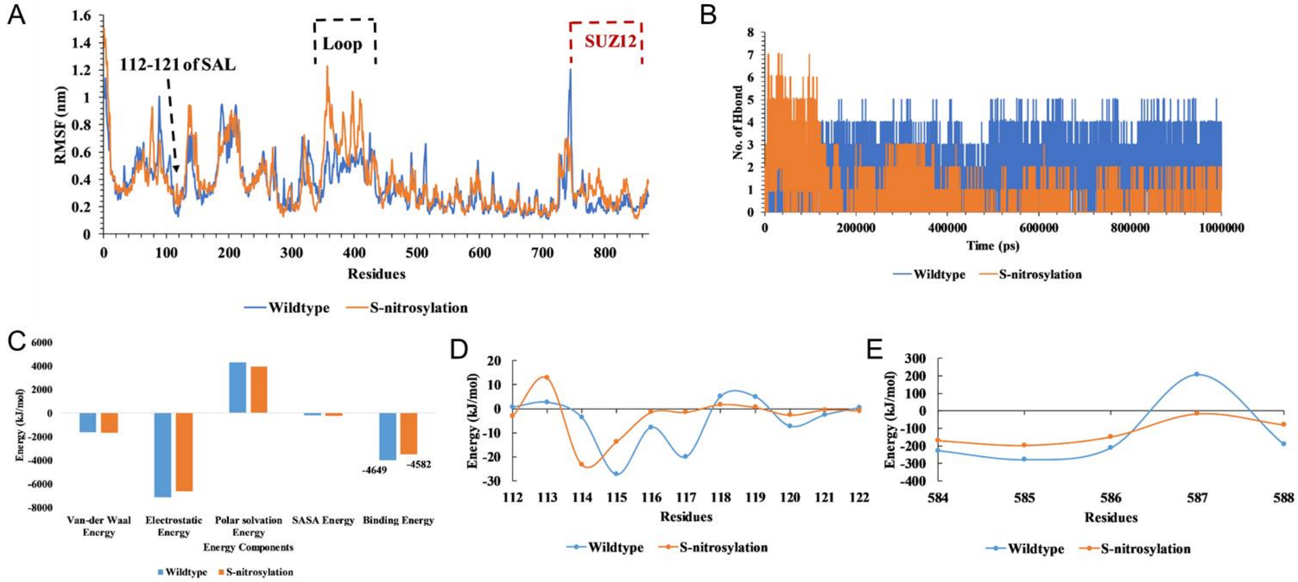


**Supplemental Figure 5. Different MD simulation based analysis of EZH2-SUZ12 complex having either EZH2 WT and EZH2 S-nitrosylated form of the protein.** (A) MD analysis using root mean square fluctuation (RMSF) to determine the flexibility of protein residues for both WT and S-nitrosylated SAL (region of EZH2) and SUZ-12-The RMSF value of both S-nitrosylated and WT residues remained relatively stable with a nearly similar trend, whereas the residues near the SAL and long loop region (residue 112-121 and 345-421) displayed significant fluctuations. (B) H-bond contact formed between residues 112-121 of SAL and 584-588 of SUZ12 over the course of 1μs MD simulation. (C) MMPBSA binding free energy. (D) Binding energy per residue of 112-121 of EZH2, (E) Binding energy per residue of 584-588 of SUZ12.
