## Supplementary Tables 1-5 for "S-nitrosylation of EZH2 at C329 and C700 interplay with PRC2 complex assembly, methyltransferase activity, and EZH2 stability to regulate endothelial functions"

**Supplementary Table 1. List of all unique proteins associated with EZH2 in untreated (control) EA.hy926 cells**

| S.NO | Description | Exp. q-value: Combined | Sum PEP Score | Coverage [%] | # Peptides | # PSMs | # Unique Peptides | # Peptides (by Search Engine): MS Amanda 2.0 | # Peptides (by Search Engine): Sequest HT |
|---|---|---|---|---|---|---|---|---|---|
| 1. | Vitamin D-binding protein OS=Homo sapiens OX=9606 GN=GC PE=1 SV=2 | 0 | 12.02 | 10 | 3 | 5 | 3 | 2 | 3 |
| 2. | Keratin, type I cuticular Ha4 OS=Homo sapiens OX=9606 GN=KRT34 PE=1 SV=2 | 0 | 7.491 | 10 | 5 | 14 | 3 | 5 | 5 |
| 3. | Fibronectin OS=Homo sapiens OX=9606 GN=FN1 PE=1 SV=5 | 0 | 5.419 | 1 | 1 | 1 | 1 | | 1 |
| 4. | Semenogelin-1 OS=Homo sapiens OX=9606 GN=SEMG1 PE=1 SV=2 | 0 | 3.294 | 7 | 2 | 4 | 2 | 2 | 2 |
| 5. | 60S ribosomal protein L23a OS=Homo sapiens OX=9606 GN=RPL23A PE=1 SV=1 | 0 | 3.018 | 13 | 2 | 4 | 2 | 2 | 2 |
| 6. | Heterogeneous nuclear ribonucleoproteins A2/B1 OS=Homo sapiens OX=9606 GN=HNRNPA2B1 PE=1 SV=2 | 0 | 2.772 | 5 | 2 | 4 | 1 | 2 | 2 |
| 7. | 60S ribosomal protein L14 | 0 | 2.617 | 6 | 1 | 2 | 1 | 1 | 1 |

| | | | | | | | | | |
|---|---|---|---|---|---|---|---|---|---|
| | OS=Homo sapiens OX=9606 GN=RPL14 PE=1 SV=4 | | | | | | | | |
| 8. | Lysosomal protective protein OS=Homo sapiens OX=9606 GN=CTSA PE=1 SV=2 | 0 | 2.438 | 3 | 1 | 2 | 1 | 1 | 1 |
| 9. | 60S ribosomal protein L22 OS=Homo sapiens OX=9606 GN=RPL22 PE=1 SV=2 | 0 | 2.428 | 10 | 1 | 2 | 1 | 1 | 1 |
| 10. | 40S ribosomal protein S15a OS=Homo sapiens OX=9606 GN=RPS15A PE=1 SV=2 | 0 | 2.277 | 18 | 2 | 4 | 2 | 2 | 2 |
| 11. | 60S ribosomal protein L8 OS=Homo sapiens OX=9606 GN=RPL8 PE=1 SV=2 | 0 | 2.186 | 4 | 1 | 2 | 1 | 1 | 1 |
| 12. | Heterogeneous nuclear ribonucleoprotein R OS=Homo sapiens OX=9606 GN=HNRNPR PE=1 SV=1 | 0 | 1.904 | 2 | 1 | 2 | 1 | 1 | 1 |
| 13. | Heterogeneous nuclear ribonucleoprotein Q OS=Homo sapiens OX=9606 GN=SYNCRIP PE=1 SV=2 | 0 | 1.904 | 2 | 1 | 2 | 1 | 1 | 1 |
| 14. | 60S ribosomal protein L7a OS=Homo sapiens OX=9606 GN=RPL7A PE=1 SV=2 | 0 | 1.85 | 5 | 1 | 2 | 1 | 1 | 1 |

| | | | | | | | | | |
|---|---|---|---|---|---|---|---|---|---|
| 15. | Semenogelin-2 OS=Homo sapiens OX=9606 GN=SEMG2 PE=1 SV=1 | 0 | 1.844 | 4 | 1 | 2 | 1 | 1 | 1 |
| 16. | Putative histone H2B type 2-C OS=Homo sapiens OX=9606 GN=H2BC20P PE=5 SV=3 | 0 | 1.69 | 5 | 1 | 2 | 1 | 1 | 1 |
| 17. | Putative histone H2B type 2-D OS=Homo sapiens OX=9606 GN=H2BC19P PE=5 SV=3 | 0 | 1.69 | 5 | 1 | 2 | 1 | 1 | 1 |
| 18. | 60S ribosomal protein L23 OS=Homo sapiens OX=9606 GN=RPL23 PE=1 SV=1 | 0.004 | 1.558 | 6 | 1 | 2 | 1 | 1 | 1 |
| 19. | Probable ATP-dependent RNA helicase DDX17 OS=Homo sapiens OX=9606 GN=DDX17 PE=1 SV=2 | 0.004 | 1.44 | 2 | 1 | 2 | 1 | 1 | 1 |
| 20. | Probable ATP-dependent RNA helicase DDX5 OS=Homo sapiens OX=9606 GN=DDX5 PE=1 SV=1 | 0.004 | 1.44 | 2 | 1 | 2 | 1 | 1 | 1 |
| 21. | Y-box-binding protein 3 OS=Homo sapiens OX=9606 GN=YBX3 PE=1 SV=4 | 0.004 | 1.38 | 2 | 1 | 2 | 1 | 1 | 1 |
| 22. | Y-box-binding protein 2 OS=Homo sapiens OX=9606 GN=YBX2 PE=1 SV=2 | 0.004 | 1.38 | 2 | 1 | 2 | 1 | 1 | 1 |

| | | | | | | | | | |
|---|---|---|---|---|---|---|---|---|---|
| 23. | Y-box-binding protein 1 OS=Homo sapiens OX=9606 GN=YBX1 PE=1 SV=3 | 0.004 | 1.38 | 2 | 1 | 2 | 1 | 1 | 1 |
| 24. | 60S ribosomal protein L35 OS=Homo sapiens OX=9606 GN=RPL35 PE=1 SV=2 | 0.004 | 1.338 | 8 | 1 | 2 | 1 | 1 | 1 |
| 25. | 40S ribosomal protein S8 OS=Homo sapiens OX=9606 GN=RPS8 PE=1 SV=2 | 0.004 | 1.325 | 5 | 1 | 1 | 1 | | 1 |
| 26. | Histone H3.3 OS=Homo sapiens OX=9606 GN=H3-3A PE=1 SV=2 | 0.004 | 1.2 | 5 | 1 | 2 | 1 | 1 | 1 |
| 27. | Histone H3.1t OS=Homo sapiens OX=9606 GN=H3-4 PE=1 SV=3 | 0.004 | 1.2 | 5 | 1 | 2 | 1 | 1 | 1 |
| 28. | Histone H3.3C OS=Homo sapiens OX=9606 GN=H3-5 PE=1 SV=3 | 0.004 | 1.2 | 5 | 1 | 2 | 1 | 1 | 1 |
| 29. | Histone H3.1 OS=Homo sapiens OX=9606 GN=H3C1 PE=1 SV=2 | 0.004 | 1.2 | 5 | 1 | 2 | 1 | 1 | 1 |
| 30. | Histone H3.2 OS=Homo sapiens OX=9606 GN=H3C15 PE=1 SV=3 | 0.004 | 1.2 | 5 | 1 | 2 | 1 | 1 | 1 |
| 31. | Proteasome subunit beta type-6 OS=Homo sapiens OX=9606 GN=PSMB6 PE=1 SV=4 | 0.004 | 1.196 | 4 | 1 | 2 | 1 | 1 | 1 |
| 32. | 60S ribosomal protein L27 OS=Homo sapiens | 0.004 | 1.189 | 7 | 1 | 2 | 1 | 1 | 1 |

| | | | | | | | | | |
|---|---|---|---|---|---|---|---|---|---|
| | OX=9606 GN=RPL27 PE=1 SV=2 | | | | | | | | |
| 33. | Olfactomedin-like protein 3 OS=Homo sapiens OX=9606 GN=OLFML3 PE=2 SV=1 | 0.004 | 1.18 | 2 | 1 | 2 | 1 | 1 | 1 |
| 34. | ADP/ATP translocase 3 OS=Homo sapiens OX=9606 GN=SLC25A6 PE=1 SV=4 | 0.007 | 1.14 | 3 | 1 | 2 | 1 | 1 | 1 |
| 35. | ADP/ATP translocase 2 OS=Homo sapiens OX=9606 GN=SLC25A5 PE=1 SV=7 | 0.007 | 1.14 | 3 | 1 | 2 | 1 | 1 | 1 |
| 36. | ADP/ATP translocase 1 OS=Homo sapiens OX=9606 GN=SLC25A4 PE=1 SV=4 | 0.007 | 1.14 | 3 | 1 | 2 | 1 | 1 | 1 |
| 37. | Phosphatidylinositol 4,5-bisphosphate 3-kinase catalytic subunit alpha isoform OS=Homo sapiens OX=9606 GN=PIK3CA PE=1 SV=2 | 0.007 | 1.08 | 1 | 1 | 1 | 1 | 1 | |
| 38. | Adenosylhomocysteinase OS=Homo sapiens OX=9606 GN=AHCY PE=1 SV=4 | 0.007 | 1.07 | 3 | 1 | 2 | 1 | 1 | 1 |
| 39. | 60S ribosomal protein L24 OS=Homo sapiens OX=9606 GN=RPL24 PE=1 SV=1 | 0.007 | 1.066 | 5 | 1 | 2 | 1 | 1 | 1 |

| | | | | | | | | | |
|---|---|---|---|---|---|---|---|---|---|
| 40. | 14-3-3 protein epsilon OS=Homo sapiens OX=9606 GN=YWHAE PE=1 SV=1 | 0.007 | 1.063 | 3 | 1 | 2 | 1 | 1 | 1 |
| 41. | Peptidyl-prolyl cis-trans isomerase A OS=Homo sapiens OX=9606 GN=PPIA PE=1 SV=2 | 0.01 | 0.96 | 5 | 1 | 2 | 1 | 1 | 1 |
| 42. | Myosin regulatory light chain 2, skeletal muscle isoform OS=Homo sapiens OX=9606 GN=MYLPF PE=1 SV=1 | 0.014 | 0.937 | 11 | 1 | 2 | 1 | 1 | 1 |
| 43. | Heterogeneous nuclear ribonucleoprotein H OS=Homo sapiens OX=9606 GN=HNRNPH1 PE=1 SV=4 | 0.014 | 0.935 | 1 | 1 | 1 | 1 | | 1 |
| 44. | Heterogeneous nuclear ribonucleoprotein H2 OS=Homo sapiens OX=9606 GN=HNRNPH2 PE=1 SV=1 | 0.014 | 0.935 | 1 | 1 | 1 | 1 | | 1 |
| 45. | Tumor necrosis factor alpha-induced protein 2 OS=Homo sapiens OX=9606 GN=TNFAIP2 PE=1 SV=2 | 0.014 | 0.924 | 2 | 1 | 2 | 1 | 1 | 1 |
| 46. | Prostaglandin-H2 D-isomerase OS=Homo sapiens OX=9606 GN=PTGDS PE=1 SV=1 | 0.013 | 0.915 | 4 | 1 | 1 | 1 | | 1 |

| | | | | | | | | | |
|---|---|---|---|---|---|---|---|---|---|
| 47. | 60S ribosomal protein L21 OS=Homo sapiens OX=9606 GN=RPL21 PE=1 SV=2 | 0.013 | 0.903 | 7 | 1 | 2 | 1 | 1 | 1 |
| 48. | Peroxisomal sarcosine oxidase OS=Homo sapiens OX=9606 GN=PIPOX PE=1 SV=2 | 0.023 | 0.858 | 8 | 1 | 1 | 1 | | 1 |

**Supplementary Table 2. List of all unique proteins associated with EZH2 in EA.hy926 cells upon GSNO treatment**

| S.No | Description | Exp. q-value: Combined | Sum PEP Score | Coverage [%] | # Peptides | # PSMs | # Unique Peptides | # Peptides (by Search Engine): MS Amanda 2.0 | # Peptides (by Search Engine): Sequest HT |
|---|---|---|---|---|---|---|---|---|---|
| 1. | Keratin, type II cytoskeletal 71 OS=Homo sapiens OX=9606 GN=KRT71 PE=1 SV=3 | 0 | 19.924 | 11 | 6 | 27 | 2 | 6 | 6 |
| 2. | Cathepsin D OS=Homo sapiens OX=9606 GN=CTSD PE=1 SV=1 | 0 | 19.908 | 23 | 9 | 19 | 9 | 8 | 9 |
| 3. | Proteasome subunit beta type-5 OS=Homo sapiens OX=9606 GN=PSMB5 PE=1 SV=3 | 0 | 9.822 | 23 | 5 | 11 | 5 | 5 | 5 |
| 4. | Heat shock protein HSP 90-alpha OS=Homo sapiens OX=9606 GN=HSP90AA1 PE=1 SV=5 | 0 | 9.085 | 8 | 5 | 11 | 2 | 5 | 4 |
| 5. | Gamma-glutamylcyclotransferase OS=Homo sapiens OX=9606 GN=GGCT PE=1 SV=1 | 0 | 8.324 | 14 | 2 | 6 | 2 | 2 | 2 |
| 6. | Apolipoprotein D OS=Homo sapiens OX=9606 GN=APOD PE=1 SV=1 | 0 | 6.496 | 20 | 4 | 11 | 4 | 4 | 4 |

| | | | | | | | | | |
|---|---|---|---|---|---|---|---|---|---|
| 7. | Neuroblast differentiation-associated protein AHNAK OS=Homo sapiens OX=9606 GN=AHNAK PE=1 SV=2 | 0 | 6.342 | 4 | 4 | 7 | 4 | 3 | 3 |
| 8. | Acid ceramidase OS=Homo sapiens OX=9606 GN=ASAH1 PE=1 SV=5 | 0 | 6.119 | 7 | 3 | 6 | 3 | 3 | 3 |
| 9. | Lysosome-associated membrane glycoprotein 1 OS=Homo sapiens OX=9606 GN=LAMP1 PE=1 SV=3 | 0 | 5.893 | 6 | 3 | 6 | 3 | 3 | 3 |
| 10. | Annexin A5 OS=Homo sapiens OX=9606 GN=ANXA5 PE=1 SV=2 | 0 | 5.796 | 8 | 2 | 4 | 2 | 2 | 2 |
| 11. | Carbonic anhydrase 1 OS=Homo sapiens OX=9606 GN=CA1 PE=1 SV=2 | 0 | 5.501 | 11 | 2 | 4 | 2 | 2 | 2 |
| 12. | Fructose-bisphosphate aldolase A OS=Homo sapiens OX=9606 GN=ALDOA PE=1 SV=2 | 0 | 5.423 | 10 | 3 | 8 | 3 | 3 | 3 |
| 13. | Uromodulin OS=Homo sapiens OX=9606 GN=UMOD PE=1 SV=1 | 0 | 5.358 | 6 | 4 | 8 | 4 | 4 | 4 |

| | | | | | | | | | |
|---|---|---|---|---|---|---|---|---|---|
| 14. | Short-chain dehydrogenase/reductase family 9C member 7 OS=Homo sapiens OX=9606 GN=SDR9C7 PE=1 SV=1 | 0 | 5.239 | 9 | 3 | 5 | 3 | 3 | 2 |
| 15. | Kallikrein-7 OS=Homo sapiens OX=9606 GN=KLK7 PE=1 SV=1 | 0 | 4.912 | 13 | 2 | 6 | 2 | 2 | 2 |
| 16. | Deleted in malignant brain tumors 1 protein OS=Homo sapiens OX=9606 GN=DMBT1 PE=1 SV=2 | 0 | 4.4 | 9 | 2 | 3 | 2 | 1 | 2 |
| 17. | Serotransferrin OS=Homo sapiens OX=9606 GN=TF PE=1 SV=3 | 0 | 4.289 | 4 | 3 | 6 | 3 | 3 | 3 |
| 18. | Gamma-glutamyl hydrolase OS=Homo sapiens OX=9606 GN=GGH PE=1 SV=2 | 0 | 4.087 | 6 | 2 | 6 | 2 | 2 | 2 |
| 19. | Bleomycin hydrolase OS=Homo sapiens OX=9606 GN=BLMH PE=1 SV=1 | 0 | 3.78 | 7 | 3 | 6 | 3 | 3 | 3 |
| 20. | Calmodulin-like protein 5 OS=Homo sapiens OX=9606 GN=CALML5 PE=1 SV=2 | 0 | 3.776 | 9 | 1 | 1 | 1 | 1 | |
| 21. | Proteasome subunit alpha type-3 OS=Homo | 0 | 3.687 | 9 | 2 | 4 | 2 | 2 | 2 |

| | | | | | | | | | |
|---|---|---|---|---|---|---|---|---|---|
| | sapiens OX=9606 GN=PSMA3 PE=1 SV=2 | | | | | | | | |
| 22. | Keratin, type I cytoskeletal 18 OS=Homo sapiens OX=9606 GN=KRT18 PE=1 SV=2 | 0 | 3.348 | 6 | 3 | 10 | 2 | 3 | 3 |
| 23. | Purine nucleoside phosphorylase OS=Homo sapiens OX=9606 GN=PNP PE=1 SV=2 | 0 | 3.195 | 7 | 2 | 5 | 2 | 2 | 2 |
| 24. | Ganglioside GM2 activator OS=Homo sapiens OX=9606 GN=GM2A PE=1 SV=4 | 0 | 3.167 | 8 | 2 | 4 | 2 | 2 | 2 |
| 25. | Alpha-1-antitrypsin OS=Homo sapiens OX=9606 GN=SERPINA1 PE=1 SV=3 | 0 | 3.144 | 5 | 2 | 4 | 2 | 2 | 2 |
| 26. | Triosephosphate isomerase OS=Homo sapiens OX=9606 GN=TPI1 PE=1 SV=4 | 0 | 2.876 | 8 | 2 | 4 | 2 | 2 | 2 |
| 27. | Galectin-3 OS=Homo sapiens OX=9606 GN=LGALS3 PE=1 SV=5 | 0 | 2.853 | 9 | 2 | 4 | 2 | 2 | 2 |
| 28. | Tropomyosin alpha-4 chain OS=Homo sapiens OX=9606 GN=TPM4 PE=1 SV=3 | 0 | 2.515 | 4 | 1 | 2 | 1 | 1 | 1 |
| 29. | Proteasome subunit alpha type-4 OS=Homo | 0 | 2.479 | 8 | 2 | 3 | 2 | 1 | 2 |

| | | | | | | | | | |
|---|---|---|---|---|---|---|---|---|---|
| | sapiens OX=9606 GN=PSMA4 PE=1 SV=1 | | | | | | | | |
| 30. | Histone H2B type 1-A OS=Homo sapiens OX=9606 GN=H2BC1 PE=1 SV=3 | 0 | 2.166 | 13 | 2 | 4 | 2 | 2 | 2 |
| 31. | Proteasome subunit alpha type-1 OS=Homo sapiens OX=9606 GN=PSMA1 PE=1 SV=1 | 0 | 2.164 | 5 | 1 | 2 | 1 | 1 | 1 |
| 32. | 40S ribosomal protein S3 OS=Homo sapiens OX=9606 GN=RPS3 PE=1 SV=2 | 0 | 2.136 | 9 | 2 | 3 | 2 | 1 | 2 |
| 33. | Carboxypeptidase A4 OS=Homo sapiens OX=9606 GN=CPA4 PE=1 SV=2 | 0 | 2.052 | 4 | 2 | 4 | 2 | 2 | 2 |
| 34. | Catenin beta-1 OS=Homo sapiens OX=9606 GN=CTNNB1 PE=1 SV=1 | 0 | 2.023 | 2 | 2 | 3 | 2 | 1 | 2 |
| 35. | Small nuclear ribonucleoprotein Sm D3 OS=Homo sapiens OX=9606 GN=SNRPD3 PE=1 SV=1 | 0 | 1.983 | 8 | 1 | 2 | 1 | 1 | 1 |
| 36. | Glyceraldehyde-3-phosphate dehydrogenase, testis-specific OS=Homo sapiens OX=9606 GN=GAPDHS PE=1 SV=2 | 0 | 1.974 | 2 | 1 | 5 | 1 | 1 | 1 |

| | | | | | | | | | |
|---|---|---|---|---|---|---|---|---|---|
| 37. | U1 small nuclear ribonucleoprotein 70 kDa OS=Homo sapiens OX=9606 GN=SNRNP70 PE=1 SV=2 | 0 | 1.926 | 5 | 2 | 4 | 2 | 2 | 2 |
| 38. | Proteasome subunit alpha-type 8 OS=Homo sapiens OX=9606 GN=PSMA8 PE=2 SV=3 | 0 | 1.92 | 4 | 1 | 2 | 1 | 1 | 1 |
| 39. | Proteasome subunit alpha type-7 OS=Homo sapiens OX=9606 GN=PSMA7 PE=1 SV=1 | 0 | 1.92 | 4 | 1 | 2 | 1 | 1 | 1 |
| 40. | Immunoglobulin lambda-like polypeptide 5 OS=Homo sapiens OX=9606 GN=IGLL5 PE=2 SV=2 | 0 | 1.765 | 7 | 1 | 2 | 1 | 1 | 1 |
| 41. | Immunoglobulin lambda constant 6 OS=Homo sapiens OX=9606 GN=IGLC6 PE=1 SV=1 | 0 | 1.765 | 14 | 1 | 2 | 1 | 1 | 1 |
| 42. | Immunoglobulin lambda constant 1 OS=Homo sapiens OX=9606 GN=IGLC1 PE=1 SV=1 | 0 | 1.765 | 14 | 1 | 2 | 1 | 1 | 1 |
| 43. | Ras-related protein Rab-10 OS=Homo sapiens OX=9606 GN=RAB10 PE=1 SV=1 | 0 | 1.729 | 6 | 1 | 2 | 1 | 1 | 1 |
| 44. | Ras-related protein Rab-1A OS=Homo | 0 | 1.729 | 5 | 1 | 2 | 1 | 1 | 1 |

| | | | | | | | | | |
|---|---|---|---|---|---|---|---|---|---|
| | sapiens OX=9606 GN=RAB1A PE=1 SV=3 | | | | | | | | |
| 45. | Ras-related protein Rab-3C OS=Homo sapiens OX=9606 GN=RAB3C PE=1 SV=1 | 0 | 1.729 | 5 | 1 | 2 | 1 | 1 | 1 |
| 46. | Ras-related protein Rab-39A OS=Homo sapiens OX=9606 GN=RAB39A PE=1 SV=2 | 0 | 1.729 | 5 | 1 | 2 | 1 | 1 | 1 |
| 47. | Ras-related protein Rab-3A OS=Homo sapiens OX=9606 GN=RAB3A PE=1 SV=1 | 0 | 1.729 | 5 | 1 | 2 | 1 | 1 | 1 |
| 48. | Ras-related protein Rab-43 OS=Homo sapiens OX=9606 GN=RAB43 PE=1 SV=1 | 0 | 1.729 | 5 | 1 | 2 | 1 | 1 | 1 |
| 49. | Ras-related protein Rab-14 OS=Homo sapiens OX=9606 GN=RAB14 PE=1 SV=4 | 0 | 1.729 | 5 | 1 | 2 | 1 | 1 | 1 |
| 50. | Ras-related protein Rab-3B OS=Homo sapiens OX=9606 GN=RAB3B PE=1 SV=2 | 0 | 1.729 | 5 | 1 | 2 | 1 | 1 | 1 |
| 51. | Ras-related protein Rab-15 OS=Homo sapiens OX=9606 GN=RAB15 PE=1 SV=1 | 0 | 1.729 | 5 | 1 | 2 | 1 | 1 | 1 |
| 52. | Ras-related protein Rab-8A OS=Homo sapiens OX=9606 GN=RAB8A PE=1 SV=1 | 0 | 1.729 | 5 | 1 | 2 | 1 | 1 | 1 |

| | | | | | | | | | |
|---|---|---|---|---|---|---|---|---|---|
| 53. | Ras-related protein Rab-6B OS=Homo sapiens OX=9606 GN=RAB6B PE=1 SV=1 | 0 | 1.729 | 5 | 1 | 2 | 1 | 1 | 1 |
| 54. | Ras-related protein Rab-12 OS=Homo sapiens OX=9606 GN=RAB12 PE=1 SV=3 | 0 | 1.729 | 5 | 1 | 2 | 1 | 1 | 1 |
| 55. | Ras-related protein Rab-4A OS=Homo sapiens OX=9606 GN=RAB4A PE=1 SV=3 | 0 | 1.729 | 5 | 1 | 2 | 1 | 1 | 1 |
| 56. | Ras-related protein Rab-33B OS=Homo sapiens OX=9606 GN=RAB33B PE=1 SV=1 | 0 | 1.729 | 5 | 1 | 2 | 1 | 1 | 1 |
| 57. | Putative Ras-related protein Rab-1C OS=Homo sapiens OX=9606 GN=RAB1C PE=5 SV=2 | 0 | 1.729 | 5 | 1 | 2 | 1 | 1 | 1 |
| 58. | Ras-related protein Rab-4B OS=Homo sapiens OX=9606 GN=RAB4B PE=1 SV=1 | 0 | 1.729 | 5 | 1 | 2 | 1 | 1 | 1 |
| 59. | Ras-related protein Rab-37 OS=Homo sapiens OX=9606 GN=RAB37 PE=1 SV=3 | 0 | 1.729 | 5 | 1 | 2 | 1 | 1 | 1 |
| 60. | Ras-related protein Rab-3D OS=Homo sapiens OX=9606 GN=RAB3D PE=1 SV=1 | 0 | 1.729 | 5 | 1 | 2 | 1 | 1 | 1 |
| 61. | Ras-related protein Rab-1B OS=Homo | 0 | 1.729 | 5 | 1 | 2 | 1 | 1 | 1 |

| | | | | | | | | | |
|---|---|---|---|---|---|---|---|---|---|
| | sapiens OX=9606 GN=RAB1B PE=1 SV=1 | | | | | | | | |
| 62. | Ras-related protein Rab-35 OS=Homo sapiens OX=9606 GN=RAB35 PE=1 SV=1 | 0 | 1.729 | 5 | 1 | 2 | 1 | 1 | 1 |
| 63. | Ras-related protein Rab-30 OS=Homo sapiens OX=9606 GN=RAB30 PE=1 SV=2 | 0 | 1.729 | 5 | 1 | 2 | 1 | 1 | 1 |
| 64. | Ras-related protein Rab-6A OS=Homo sapiens OX=9606 GN=RAB6A PE=1 SV=3 | 0 | 1.729 | 5 | 1 | 2 | 1 | 1 | 1 |
| 65. | Ras-related protein Rab-39B OS=Homo sapiens OX=9606 GN=RAB39B PE=1 SV=1 | 0 | 1.729 | 5 | 1 | 2 | 1 | 1 | 1 |
| 66. | Ras-related protein Rab-8B OS=Homo sapiens OX=9606 GN=RAB8B PE=1 SV=2 | 0 | 1.729 | 5 | 1 | 2 | 1 | 1 | 1 |
| 67. | Immunoglobulin J chain OS=Homo sapiens OX=9606 GN=JCHAIN PE=1 SV=4 | 0 | 1.662 | 8 | 1 | 2 | 1 | 1 | 1 |
| 68. | Cytosol aminopeptidase OS=Homo sapiens OX=9606 GN=LAP3 PE=1 SV=3 | 0 | 1.621 | 1 | 1 | 2 | 1 | 1 | 1 |
| 69. | Retinoid-inducible serine carboxypeptidase OS=Homo sapiens OX=9606 | 0 | 1.618 | 2 | 1 | 2 | 1 | 1 | 1 |

| | | | | | | | | | |
|---|---|---|---|---|---|---|---|---|---|
| | GN=SCPEP1 PE=1 SV=1 | | | | | | | | |
| 70. | Protein S100-A6 OS=Homo sapiens OX=9606 GN=S100A6 PE=1 SV=1 | 0 | 1.604 | 9 | 1 | 2 | 1 | 1 | 1 |
| 71. | Cystatin-M OS=Homo sapiens OX=9606 GN=CST6 PE=1 SV=1 | 0 | 1.579 | 7 | 1 | 2 | 1 | 1 | 1 |
| 72. | Serpin B7 OS=Homo sapiens OX=9606 GN=SERPINB7 PE=1 SV=1 | 0 | 1.575 | 3 | 1 | 2 | 1 | 1 | 1 |
| 73. | 60S ribosomal protein L36 OS=Homo sapiens OX=9606 GN=RPL36 PE=1 SV=3 | 0 | 1.541 | 9 | 1 | 2 | 1 | 1 | 1 |
| 74. | Calpain-1 catalytic subunit OS=Homo sapiens OX=9606 GN=CAPN1 PE=1 SV=1 | 0 | 1.515 | 1 | 1 | 2 | 1 | 1 | 1 |
| 75. | Destrin OS=Homo sapiens OX=9606 GN=DSTN PE=1 SV=3 | 0 | 1.447 | 4 | 1 | 2 | 1 | 1 | 1 |
| 76. | Proteasome subunit beta type-7 OS=Homo sapiens OX=9606 GN=PSMB7 PE=1 SV=1 | 0 | 1.426 | 3 | 1 | 2 | 1 | 1 | 1 |
| 77. | 60S ribosomal protein L13 OS=Homo sapiens OX=9606 GN=RPL13 PE=1 SV=4 | 0 | 1.39 | 6 | 1 | 2 | 1 | 1 | 1 |

| | | | | | | | | | |
|---|---|---|---|---|---|---|---|---|---|
| 78. | Transmembrane glycoprotein NMB OS=Homo sapiens OX=9606 GN=GPNMB PE=1 SV=2 | 0 | 1.364 | 3 | 1 | 2 | 1 | 1 | 1 |
| 79. | Apolipoprotein A-II OS=Homo sapiens OX=9606 GN=APOA2 PE=1 SV=1 | 0 | 1.344 | 9 | 1 | 2 | 1 | 1 | 1 |
| 80. | Speckle targeted PIP5K1A-regulated poly(A) polymerase OS=Homo sapiens OX=9606 GN=TUT1 PE=1 SV=2 | 0 | 1.341 | 1 | 1 | 1 | 1 | 1 | |
| 81. | Leucine-rich alpha-2-glycoprotein OS=Homo sapiens OX=9606 GN=LRG1 PE=1 SV=2 | 0 | 1.322 | 3 | 1 | 2 | 1 | 1 | 1 |
| 82. | Loricrin OS=Homo sapiens OX=9606 GN=LORICRIN PE=1 SV=2 | 0 | 1.316 | 3 | 1 | 4 | 1 | 1 | 1 |
| 83. | Malate dehydrogenase, mitochondrial OS=Homo sapiens OX=9606 GN=MDH2 PE=1 SV=3 | 0 | 1.3 | 3 | 1 | 2 | 1 | 1 | 1 |
| 84. | Translocator protein OS=Homo sapiens OX=9606 GN=TSPO PE=1 SV=3 | 0 | 1.246 | 5 | 1 | 2 | 1 | 1 | 1 |
| 85. | Testis-specific Y-encoded-like protein 2 OS=Homo sapiens OX=9606 | 0 | 1.214 | 1 | 1 | 2 | 1 | 1 | 1 |

| | | | | | | | | | |
|---|---|---|---|---|---|---|---|---|---|
| | GN=TSPYL2 PE=1 SV=1 | | | | | | | | |
| 86. | Heterogeneous nuclear ribonucleoprotein M OS=Homo sapiens OX=9606 GN=HNRNPM PE=1 SV=3 | 0 | 1.211 | 2 | 1 | 2 | 1 | 1 | 1 |
| 87. | Proteasome subunit alpha type-5 OS=Homo sapiens OX=9606 GN=PSMA5 PE=1 SV=3 | 0.003 | 1.208 | 3 | 1 | 1 | 1 | 1 | |
| 88. | Insulin-degrading enzyme OS=Homo sapiens OX=9606 GN=IDE PE=1 SV=4 | 0.006 | 1.194 | 1 | 1 | 2 | 1 | 1 | 1 |
| 89. | Very-long-chain enoyl-CoA reductase OS=Homo sapiens OX=9606 GN=TECR PE=1 SV=1 | 0.009 | 1.174 | 3 | 1 | 2 | 1 | 1 | 1 |
| 90. | Sesquipedalian-1 OS=Homo sapiens OX=9606 GN=PHETA1 PE=1 SV=1 | 0.009 | 1.156 | 4 | 1 | 1 | 1 | | 1 |
| 91. | 60S ribosomal protein L28 OS=Homo sapiens OX=9606 GN=RPL28 PE=1 SV=3 | 0.009 | 1.149 | 5 | 1 | 2 | 1 | 1 | 1 |
| 92. | Histone H2B type 2-E1 OS=Homo sapiens OX=9606 GN=H2BE1 PE=3 SV=1 | 0.011 | 1.113 | 7 | 1 | 2 | 1 | 1 | 1 |
| 93. | Epiplakin OS=Homo sapiens OX=9606 | 0.011 | 1.11 | 1 | 1 | 2 | 1 | 1 | 1 |

| | | | | | | | | | |
|---|---|---|---|---|---|---|---|---|---|
| | GN=EPPK1 PE=1 SV=3 | | | | | | | | |
| 94. | Plectin OS=Homo sapiens OX=9606 GN=PLEC PE=1 SV=3 | 0.011 | 1.11 | 0 | 1 | 2 | 1 | 1 | 1 |
| 95. | STE20-like serine/threonine-protein kinase OS=Homo sapiens OX=9606 GN=SLK PE=1 SV=1 | 0.011 | 1.106 | 1 | 1 | 4 | 1 | 1 | 1 |
| 96. | Cytospin-B OS=Homo sapiens OX=9606 GN=SPECC1 PE=1 SV=1 | 0.011 | 1.106 | 1 | 1 | 4 | 1 | 1 | 1 |
| 97. | Chromobox protein homolog 6 OS=Homo sapiens OX=9606 GN=CBX6 PE=1 SV=1 | 0.014 | 1.07 | 2 | 1 | 2 | 1 | 1 | 1 |
| 98. | Chromobox protein homolog 8 OS=Homo sapiens OX=9606 GN=CBX8 PE=1 SV=3 | 0.014 | 1.07 | 2 | 1 | 2 | 1 | 1 | 1 |
| 99. | Rho guanine nucleotide exchange factor 40 OS=Homo sapiens OX=9606 GN=ARHGEF40 PE=1 SV=3 | 0.014 | 1.065 | 1 | 1 | 2 | 1 | 1 | 1 |
| 100. | Myosin-10 OS=Homo sapiens OX=9606 GN=MYH10 PE=1 SV=3 | 0.014 | 1.053 | 1 | 1 | 1 | 1 | 1 | |
| 101. | Vinculin OS=Homo sapiens OX=9606 GN=VCL PE=1 SV=4 | 0.014 | 1.048 | 1 | 1 | 1 | 1 | | 1 |

| | | | | | | | | | |
|---|---|---|---|---|---|---|---|---|---|
| 102. | Heterogeneous nuclear ribonucleoprotein A1-like 2 OS=Homo sapiens OX=9606 GN=HNRNPA1L2 PE=2 SV=2 | 0.014 | 1.045 | 4 | 1 | 1 | 1 | | 1 |
| 103. | Lamin-B1 OS=Homo sapiens OX=9606 GN=LMNB1 PE=1 SV=2 | 0.014 | 1.043 | 1 | 1 | 2 | 1 | 1 | 1 |
| 104. | Lamin-B2 OS=Homo sapiens OX=9606 GN=LMNB2 PE=1 SV=4 | 0.014 | 1.043 | 1 | 1 | 2 | 1 | 1 | 1 |
| 105. | Heat shock 70 kDa protein 6 OS=Homo sapiens OX=9606 GN=HSPA6 PE=1 SV=2 | 0.014 | 1.032 | 1 | 1 | 2 | 1 | 1 | 1 |
| 106. | Heat shock 70 kDa protein 1B OS=Homo sapiens OX=9606 GN=HSPA1B PE=1 SV=1 | 0.014 | 1.032 | 1 | 1 | 2 | 1 | 1 | 1 |
| 107. | Heat shock 70 kDa protein 1-like OS=Homo sapiens OX=9606 GN=HSPA1L PE=1 SV=2 | 0.014 | 1.032 | 1 | 1 | 2 | 1 | 1 | 1 |
| 108. | Toll-interacting protein OS=Homo sapiens OX=9606 GN=TOLLIP PE=1 SV=1 | 0.013 | 1.025 | 3 | 1 | 2 | 1 | 1 | 1 |
| 109. | Reticulon-4 OS=Homo sapiens | 0.013 | 1.025 | 1 | 1 | 1 | 1 | | 1 |

| | | | | | | | | | |
|---|---|---|---|---|---|---|---|---|---|
| | OX=9606 GN=RTN4 PE=1 SV=2 | | | | | | | | |
| 110. | Ergosterol biosynthetic protein 28 homolog OS=Homo sapiens OX=9606 GN=ERG28 PE=1 SV=1 | 0.021 | 1.005 | 6 | 1 | 2 | 1 | 1 | 1 |
| 111. | Histidine--tRNA ligase, cytoplasmic OS=Homo sapiens OX=9606 GN=HARS1 PE=1 SV=2 | 0.021 | 1.003 | 2 | 1 | 1 | 1 | | 1 |
| 112. | Stress-70 protein, mitochondrial OS=Homo sapiens OX=9606 GN=HSPA9 PE=1 SV=2 | 0.021 | 1.003 | 1 | 1 | 3 | 1 | 1 | 1 |
| 113. | Exocyst complex component 4 OS=Homo sapiens OX=9606 GN=EXOC4 PE=1 SV=1 | 0.021 | 0.998 | 2 | 1 | 3 | 1 | 1 | 1 |
| 114. | Cystatin-SA OS=Homo sapiens OX=9606 GN=CST2 PE=1 SV=1 | 0.024 | 0.963 | 5 | 1 | 2 | 1 | 1 | 1 |
| 115. | Leucine-rich repeat-containing protein 15 OS=Homo sapiens OX=9606 GN=LRRC15 PE=2 SV=2 | 0.029 | 0.947 | 2 | 1 | 2 | 1 | 1 | 1 |
| 116. | Tubulin alpha chain-like 3 OS=Homo sapiens OX=9606 | 0.029 | 0.942 | 2 | 1 | 2 | 1 | 1 | 1 |

| | | | | | | | | | |
|---|---|---|---|---|---|---|---|---|---|
| | GN=TUBAL3 PE=1 SV=2 | | | | | | | | |
| 117. | Tubulin alpha-8 chain OS=Homo sapiens OX=9606 GN=TUBA8 PE=1 SV=1 | 0.029 | 0.942 | 2 | 1 | 2 | 1 | 1 | 1 |
| 118. | Tubulin alpha-4A chain OS=Homo sapiens OX=9606 GN=TUBA4A PE=1 SV=1 | 0.029 | 0.942 | 2 | 1 | 2 | 1 | 1 | 1 |
| 119. | Uncharacterized protein CCDC197 OS=Homo sapiens OX=9606 GN=CCDC197 PE=1 SV=2 | 0.031 | 0.914 | 7 | 1 | 1 | 1 | | 1 |
| 120. | Ras-related C3 botulinum toxin substrate 3 OS=Homo sapiens OX=9606 GN=RAC3 PE=1 SV=1 | 0.031 | 0.903 | 4 | 1 | 2 | 1 | 1 | 1 |
| 121. | Ras-related C3 botulinum toxin substrate 2 OS=Homo sapiens OX=9606 GN=RAC2 PE=1 SV=1 | 0.031 | 0.903 | 4 | 1 | 2 | 1 | 1 | 1 |
| 122. | Ras-related C3 botulinum toxin substrate 1 OS=Homo sapiens OX=9606 GN=RAC1 PE=1 SV=1 | 0.031 | 0.903 | 4 | 1 | 2 | 1 | 1 | 1 |
| 123. | Prosaposin OS=Homo sapiens OX=9606 GN=PSAP PE=1 SV=2 | 0.031 | 0.889 | 2 | 1 | 1 | 1 | | 1 |
| 124. | Transcription factor AP-2-delta OS=Homo sapiens | 0.031 | 0.886 | 2 | 1 | 1 | 1 | | 1 |

| | | | | | | | | | |
|---|---|---|---|---|---|---|---|---|---|
| | OX=9606 GN=TFAP2D PE=1 SV=1 | | | | | | | | |
| 125. | 40S ribosomal protein S4, X isoform OS=Homo sapiens OX=9606 GN=RPS4X PE=1 SV=2 | 0.031 | 0.882 | 3 | 1 | 1 | 1 | | 1 |
| 126. | Serpin B10 OS=Homo sapiens OX=9606 GN=SERPINB10 PE=1 SV=1 | 0.031 | 0.867 | 3 | 1 | 1 | 1 | | 1 |
| 127. | H(+)/Cl(-) exchange transporter 3 OS=Homo sapiens OX=9606 GN=CLCN3 PE=1 SV=2 | 0.03 | 0.85 | 1 | 1 | 2 | 1 | 1 | 1 |
| 128. | Fibrinogen gamma chain OS=Homo sapiens OX=9606 GN=FGG PE=1 SV=3 | 0.03 | 0.841 | 3 | 1 | 2 | 1 | 1 | 1 |

**Supplementary Table 3.** Contribution of individual interaction components and binding free energy for the EZH2-SUZ12 Complex.

| Compounds | Van-der Waal energy (kJ/mol) | Electrostatic energy (kJ/mol) | Polar solvation energy (kJ/mol) | SASA energy (kJ/mol) | Binding energy (kJ/mol) |
|---|---|---|---|---|---|
| Wildtype | -1630 ± 62 | -7099 ± 618 | 4275 ± 332 | -196 ± 10 | -4649 ± 454 |
| S-nitrosylation | -1657 ± 76 | -6642 ± 706 | 3923 ± 351 | -206.485 ± 7 | -4583 ± 453 |

### Supplementary Table 4. List primers for qPCR analysis

| Primer Name | Sequence (5'->3') |
|---|---|
| KDR Forward | CCTCCTTCTCTAGACAGGCG |
| KDR Reverse | CCTCTGTCCCCTGCAAGTAA |
| TBX20 Forward | ACAGCCTCATTGCTCAACCT |
| TBX20 Reverse | GCTCTCCACACTTTCCCTCT |
| VEGFAa Forward | GGCCAGCACATAGGAGAGAT |
| VEGFAa Reverse | ACGCTCCAGGACTTATACCG |
| MMP2 Forward | CTACTGAGTGGCCGTGTTTG |
| MMP2 Reverse | TCCCTGAGGTTCTCTTGCTG |
| TIE 1 Forward | CAGCCTCTACCCTTAGCTCC |
| TIE 1 Reverse | AAAGGCCGAAGTCTGCAATC |
| TEK Forward | TGGACAAGAGGGATGCAAGT |
| TEK Reverse | TGCCTTCTCTCTCACACTGG |
| FGF2 Forward | AGTCTTCGCCAGGTCATTGA |
| FGF2 Reverse | CCTGAGTATTCGGCAACAGC |
| Angiopoietin 2Forward | GTGTCCTCTTCCACCACAGA |
| Angiopoietin 2 Reverse | TCAGCCTCGGGTTCATCTTT |

### Supplementary Table 5. List primers for inserting point mutation at specific locations

| Primer Name | Sequence (5'>3') |
|---|---|
| EZH2 -260<br>Forward<br>Reverse | <br>TCCTCCTGAAAGTACCCCCAACATAGATGG<br>AGTGCGCCTGGGAGCTGC |
| EZH2 329<br>Forward<br>Reverse | <br>TGGACCACAGAGTTACCAGCATT<br>CAAGGTTTGTTGTCTAGAGC |
| EZH2 720<br>Forward<br>Reverse | <br>AAATCCAAACAGCTATGCAAAAG<br>ACCGAATGATTTGCAAAAC |
